# Hydrophobic interactions of FG-nucleoporins are required for dilating nuclear membrane pores into selective transport channels after mitosis

**DOI:** 10.1101/2025.09.08.674396

**Authors:** Wanlu Zhang, Andrew P. Latham, Paolo Ronchi, Sebastian Schnorrenberg, Jean-Karim Hériché, Ziqiang Huang, M. Julius Hossain, Natalia Rosalia Morero, Hannah Pflaumer, Merle Hantsche-Grininger, Yannick Schwab, Andrej Sali, Jan Ellenberg

## Abstract

Nuclear envelope (NE) reformation after mitosis is essential for daughter cell viability and requires tightly coordinated nuclear pore complex (NPC) assembly and nuclear membrane reformation. To reveal how these processes are mechanistically linked, we combined acute molecule perturbations in live cells with correlative 3D electron tomography or MINFLUX super-resolution microscopy. We show that degrading Nup62 during mitosis arrests NPC assembly at an intermediate step with smaller membrane pores and removes the whole central transport channel. Molecular dynamics simulations predicted that 32 copies of the central channel subcomplex, recruited into the previously unoccupied pore center, can self-associate via hydrophobic interactions to occupy the volume required for full pore size and exert an outward pushing force; indeed, disrupting these interactions during NPC assembly blocked pore dilation. Later in mitotic exit, perturbed cells exhibited impaired nuclear import, smaller nuclei, and looser NE spacing. Acute inhibition of nuclear import recapitulated these NE defects without affecting NPC assembly. Together, our findings reveal a new, two-step molecular mechanism linking NPC assembly and NE reformation. First, hydrophobic FG-nucleoporins dilate the assembling nuclear pore to its full width by forming the central transport channel, which then allows nuclear import-driven nuclear expansion leading to tight, regular NE membrane spacing.

## Introduction

The barrier of the largest cellular compartment, the nucleus, is composed of a double-layered membrane system, the nuclear envelope (NE), perforated by large proteinaceous transport channels, the nuclear pore complexes (NPCs). In metazoan cells, this barrier disassembles at mitotic entry to allow chromosome segregation and must be rapidly and properly reassembled during mitotic exit to restore nuclear architecture and cellular functions. This reassembly occurs remarkably fast, completing within ∼20 min after anaphase onset (AO) ^1–9^. It involves two tightly coordinated processes: rapid recruitment of endoplasmic reticulum (ER)-derived membranes to coat segregated chromosomes, forming the new nuclear membranes; and the simultaneous assembly of thousands of NPCs into small fenestrae of the reforming nuclear membranes to restore nuclear transport competence ^1,3,4,10^. Failure of either process can lead to severe cellular defects, including impaired nuclear transport, aberrant nuclear architecture, formation of micronuclei, and increased risk of genomic instability ^11–16^.

NPCs are the largest non-polymeric protein assemblies in eukaryotic cells, with a molecular mass ranging from 60 to 120 MDa. A single NPC is built from multiple copies (8-64) of ∼30 different proteins (termed nucleoporins, Nups), resulting a total of ∼500-1000 proteins, organized into a rotationally symmetrical, octameric complex. The structural complexity of the complex makes its assembly a formidable cellular task, and correspondingly, a major challenge to characterize mechanistically, particularly during the brief window of rapid NE reformation at the mitotic exit. In human cells, this relatively synchronous wave of assembling thousands of NPCs, known as postmitotic NPC assembly, begins ∼5 min after AO in the non-core region of the reforming NE ^1,3^. There, Nups are sequentially incorporated into pre-pore structures, which gradually dilate to form thousands of mature, transport-competent NPCs within about 10 min ^1,3,17,18^. Our recent study provided the first spatiotemporal model of postmitotic NPC assembly, which predicts key intermediate structures and transition states ^4,10^. The model suggests that the recruitment of the central FG-Nup Nup62 coincides with pore dilation, a crucial structural transition during NPC maturation. However, whether and how Nup62 contributes to this transition remains unresolved.

Nup62 participates in two distinct subcomplexes within the NPC ^19–23^. On the cytoplasmic side, 16 copies of Nup62 form heterotrimeric subcomplexes with Nup214 and Nup88 through coiled-coil domain interactions ^20,23^, functioning as part of the nuclear export machinery ^24–26^. In the central channel, 32 copies of Nup62 associate with Nup58 and Nup54 through similar coiled-coil domain interactions ^22^, while the intrinsically disordered regions of the subcomplexes are enriched in FG repeats, which mediate the active transport together with nuclear transport receptors ^27–29^. Hydrophobic FG repeats can undergo liquid-liquid phase separation *in vitro* ^30–36^, and similar cohesive interactions are important for establishing NPC cargo selectivity ^37–40^. In contrast, the role of interactions between the FG repeats of FG-Nups in the NPC assembly process remains to be characterized.

During mitotic exit, the concurrence of NPC assembly and NE reformation suggests a mechanistic interdependence between these two processes. Correlative electron microscopy has revealed that the reforming NE undergoes a gradual structural transition during this period, from a loosely spaced double membrane to a tightly aligned and parallel structure, shortly after NPC maturation is complete ^3^. This observation raises a key mechanistic question: could NPC maturation contribute to reshaping the NE architecture? Intriguingly, defects in NPC assembly are associated with abnormal nuclear reformation, including defects in nuclear size, altered chromatin organization, and increased genomic instability ^12,14–16,41^, underscoring the critical role of functionally assembled NPCs in reestablishing nuclear integrity. However, the molecular principles linking NPC assembly and NE membrane architecture remain poorly understood.

To address these questions, we employed an integrated approach, including acute molecule perturbations, single-cell live imaging correlated with high-resolution 3D electron tomography (ET), super-resolution imaging with 3D minimal fluorescence photon fluxes microscopy (MINFLUX), and molecular simulations. We demonstrate that Nup62 is essential for pore dilation and NE flattening during mitotic exit, acting through hydrophobic interactions with disordered regions of the central FG-Nups (Nup62, Nup58, and Nup54). These interactions in the central channel drive pore dilation and modulate the NE architecture, mechanistically linking NPC maturation and NE remodeling during mitotic exit.

## Results

### Nup62 depletion during mitosis reduces the diameter of assembling pores and increases nuclear membrane spacing

Our spatiotemporal integrative model of postmitotic NPC assembly ^4,10^ predicted that Nup62 is recruited to the assembling pore before it dilates to its full diameter. This prediction raised a key mechanistic question: does Nup62 actively drive pore dilation, or is its recruitment merely coincidental?

To address this question, we established a live cell degradation strategy to deplete Nup62 rapidly during mitosis (Fig. 1A), avoiding the nonspecific effects of removing this essential protein for longer times from cells. Using our fast CRISPR-Cas9 editing platform ^42^, we homozygously tagged all endogenous Nup62 alleles in HeLa cells with a bifunctional mEGFP-FKBP12^F36V^ degron tag (Fig. 1 Suppl. 1A and B) ^43,44^, enabling rapid and selective degradation of ∼90% of Nup62 within 1 hour (Fig. 1 Suppl. 1C-F).

**Figure 1.**
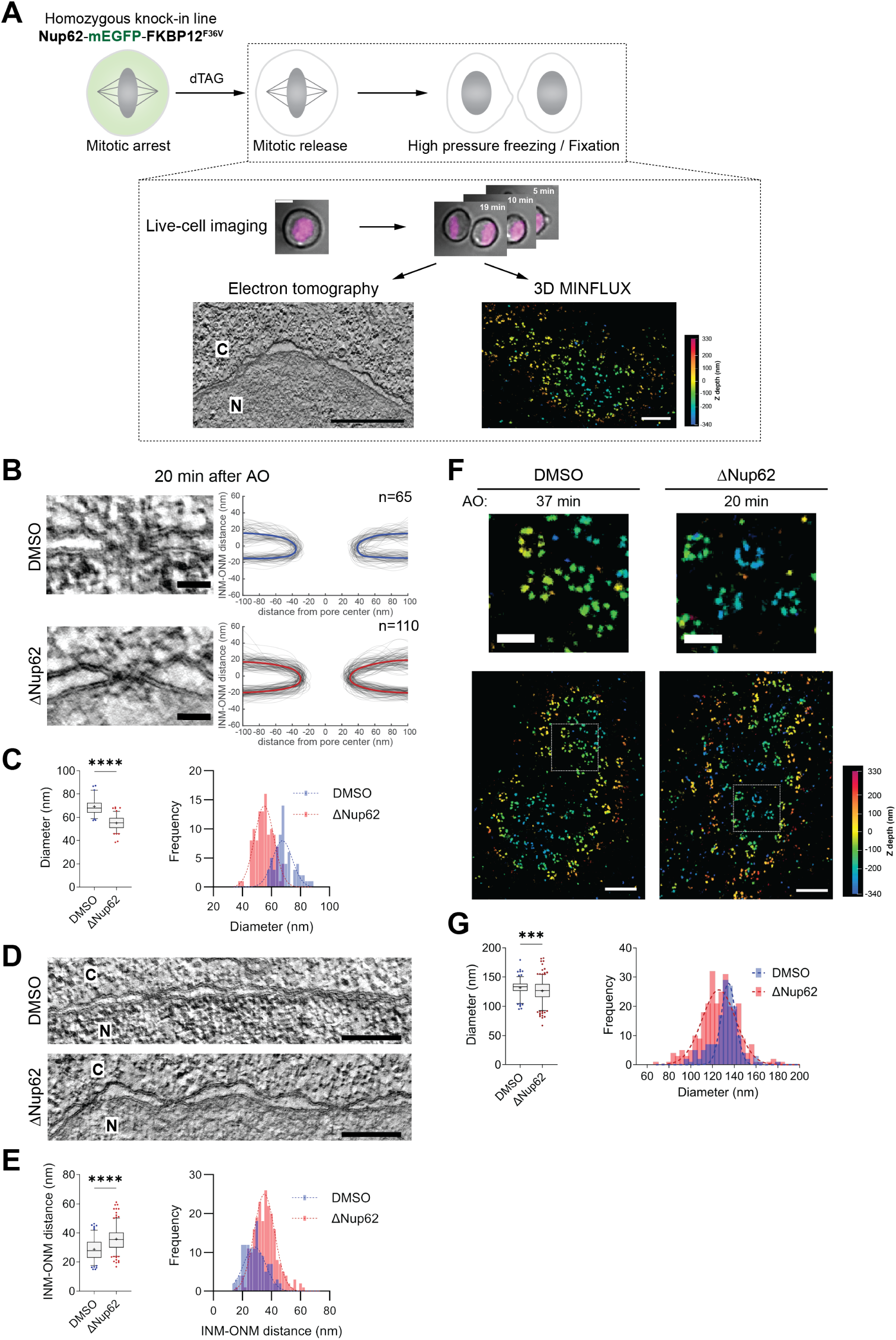
Nup62 depletion during mitosis reduces assembled pore diameter and increases NE spacing. **(A)** Experimental scheme illustrating acute depletion of endogenous Nup62-mEGFP-FKBP^F36V^ during mitosis, correlating live imaging with electron tomography or 3D MINFLUX imaging. HK Nup62-mEGFP-FKBP^F36V^ cells were arrested in prometaphase and treated with degradation compounds dTAG (250 nM dTAG-13 and 500 nM dTAG^V^-1) for 90 min before release into mitotic exit; DMSO-treated cells served as controls. Mitotic progression was monitored every 1 min by light microscopy using live DNA dyes (shown in magenta, upper panel). Cells were then either high-pressure frozen or formaldehyde-fixed at defined time points after anaphase onset (AO). Lower-left panel: transmission electron microscope (TEM) imaging of the NE from relocated cells of interest. N, nucleus; C, cytoplasm. Lower-right panel: 3D MINFLUX imaging of anti-Elys-labeled cells using the DNA-PAINT approach. Scale bars, upper panel: light microscopy, 10 μm; lower-left panel: NE tomography, 500 nm; lower-right panel: 3D MINFLUX, 500 nm. **(B)** Left, tomographic slices showing cross-section views of nuclear pores assembled under DMSO or dTAG (ΔNup62) treatment at 20 min after AO. Right, membrane profiles of all measured pores are displayed, with mean profiles highlighted in bold. Scale bars, 50 nm. **(C)** Quantification of the NPC diameter based on (B). Left, box-and-whisker plot showing median, mean (“+”), 5–95 percentiles, and outliers (scatter). Right, histogram with Gaussian fits (dashed curves). Sample sizes as indicated in (B). (**D)** Representative tomographic slices of the NE from cells treated with DMSO or dTAG (ΔNup62) at 20 min after AO. N, nucleus; C, cytoplasm. Scale bars, 200 nm. **(E)** Quantification of the INM-ONM distance from cells in (D). Left, box-and-whisker plot showing median, mean (“+”), 5–95 percentiles, and outliers (scatter). Right, histogram with Gaussian fits (dashed curves). **(F)** Representative 3D MINFLUX images (anti-Elys) of cells treated with DMSO or dTAG (ΔNup62) at the indicated time points after AO. Scale bars: upper panel, 200 nm; lower panel, 500 nm. **(G)** Quantification of the NPC diameter based on (F) from cells dividing between 20 and 60 min after AO. Left, box-and-whisker plot showing median, mean (“+”), 5–95 percentiles, and outliers (scatter). Right, histogram with Gaussian fits (dashed curves). Sample sizes: DMSO, n=150; ΔNup62, n=276. Statistical significance applies to all panels in this figure: *, P ≤ 0.05; **, P ≤ 0.01; ***, P ≤ 0.001; ****, P ≤ 0.0001; ns, not significant.

To determine the structural consequences of Nup62 depletion for NPC assembly and NE reformation, we performed correlative high-resolution transmission electron tomography (ET) of cells exiting mitosis after Nup62 degradation. Live-cell imaging at 1 min intervals after AO, prior to EM fixation, enabled precise temporal staging of mitotic progression (Fig. 1A and Fig. 1 Suppl. 2A). The electron tomograms revealed a clear defect of pore dilation in Nup62-depleted cells at 20 min after AO, when postmitotic NPC assembly is typically complete ^3,4^. In these cells, tracing of membrane profiles showed pores arrested at an intermediate state with a significantly reduced average diameter of ∼55 nm, compared to ∼70 nm in control cells with normal levels of Nup62 (Fig. 1B and C; Fig. 1 Suppl. 2B). This failure in pore dilation also persisted at later timepoints (Fig. 1 Suppl. 2C and E; Fig. 1 Suppl. 3A), showing that NPC assembly is initiated but fails to progress to full dilation in the absence of Nup62. In addition to narrower pores, Nup62-depleted cells also displayed a significant increase in the spacing between the inner and outer nuclear membranes, in contrast to control cells (Fig. 1D and E). This wider and much less regular double-membrane spacing also persisted through later timepoints after mitotic exit (Fig. 1 Suppl. 2D and F; Fig. 1 Suppl. 3B).

To probe if the partially dilated pores in Nup62-depleted cells also had a changed molecular architecture, we complemented electron tomography with correlative 3D MINFLUX super-resolution microscopy after live cell staging, using the nuclear outer ring component Elys as a marker of the structural scaffold of the NPC (Fig. 1A) ^45^. Ring-like NPC structures could clearly be visualized by MINFLUX in postmitotic cells, allowing us to determine the diameters of single NPC assembly intermediates (Methods). In Nup62-depleted cells, the diameter of Elys-labeled rings was significantly reduced (average ∼126 nm) compared to DMSO-treated controls (∼133 nm) (Fig. 1F and G), a difference that also persisted at later time points (Fig. 1 Suppl. 4A). The reduction of the outer ring diameter as labeled by Elys was consistent with the narrowing of the central membrane pore observed by electron tomography based on membrane tip-to-tip measurements at their closest point, i.e., the neck of the pore (Fig. 1B and C; Fig. 1 Suppl. 2C and E; Fig. 1 Suppl. 3A). The reduction of the peripheral outer ring diameter is smaller than that of the pore neck (7 compared to 15 nm), likely reflecting the plasticity of the NPC structure ^46,47^.

Together, our findings demonstrate that Nup62 plays an active and essential role during mitotic exit. Specifically, Nup62 is essential for full dilation of the central NPC membrane pore, maturation of the peripheral ring, and for establishing a tight and even spacing between the inner and outer nuclear membrane.

### Nup62-depleted NPC assembly intermediates lack key subcomplexes

Because Nup62 recruitment occurs at the midpoint of NPC assembly ^4^, we next wanted to probe if its acute depletion in mitosis would affect the molecular composition of assembly intermediates. Thus, to systematically analyze the presence of different Nups in assembly intermediates in Nup62-depleted cells, we used either live-cell imaging to directly follow mScarlet-tagged Nups transiently expressed in cells, or again a correlative approach combining live imaging with quantitative ratiometric immunofluorescence to map mitotic staging (Fig. 2A). As a control, we used the nuclear basket protein Nup153, which is recruited at the beginning of NPC assembly ^4^ independently from the presence of Nup62 (Fig. 2 Suppl. 1A).

**Figure 2.**
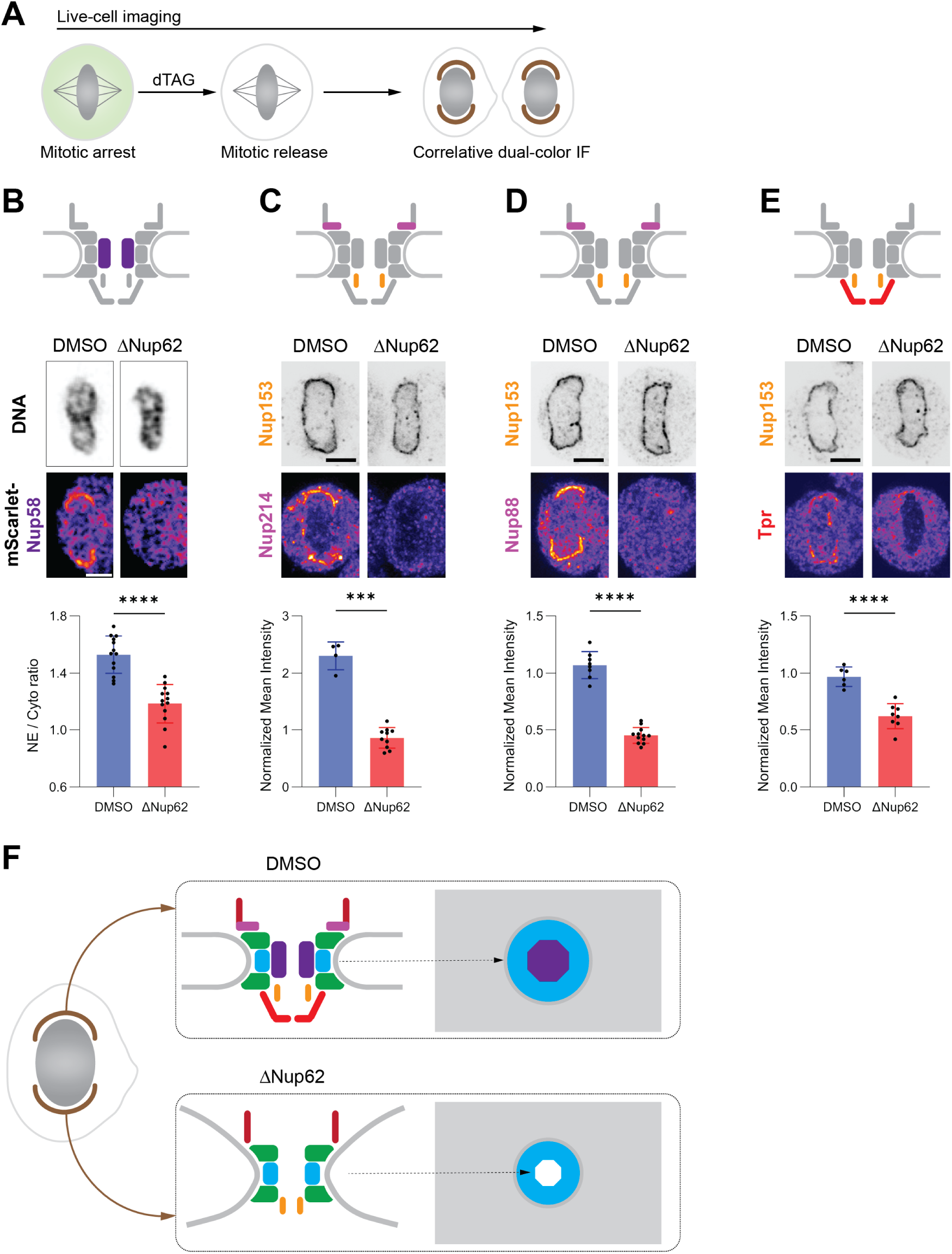
Loss of Nup62 results in incomplete NPC assembly during mitotic exit. **(A)** Experimental scheme combining acute depletion of Nup62-mEGFP-FKBP^F36V^ with correlative immunofluorescence. HK Nup62-mEGFP-FKBP^F36V^ cells were arrested in prometaphase and treated with degradation compounds dTAG (250 nM dTAG-13 and 500 nM dTAG^V^-1) for 90 min before release into mitotic exit; DMSO-treated cells served as controls. Mitotic progression was monitored every 1 min by light microscopy using live DNA dyes. Cells were then fixed at different time points after AO and relocated for immunofluorescence imaging after antibody labeling. **(B)** Representative live-cell images of HK Nup62-mEGFP-FKBP^F36V^ cells expressing mScarlet-Nup58 (middle panel), treated with DMSO or dTAG (ΔNup62) at 20 min after AO, Scale bar, 5 μm. Upper panel: NPC schematic indicating the location of Nup58. Lower panel: NE-to-cytoplasm mean intensity ratio of mScarlet-Nup58 in dividing cells at 20 min after AO. **(C-E)** Representative immunofluorescence images (middle panel) of HK Nup62-mEGFP-FKBP^F36V^ cells treated with DMSO or dTAG (ΔNup62) at 20 min after AO, dually stained with anti-Nup153 and anti-Nup214 (B), anti-Nup88 (C), or anti-Tpr (D). Scale bars, 5 μm. Upper panel: NPC schematic indicating the position of each labeled Nup within the complex. Lower panel: normalized mean intensity (calculation detailed in Fig. 2 Suppl. 1) from non-core NE regions of dividing cells at 20 ± 1 min after AO. **(F)** Schematic representation of NPC structure assembled in the non-core NE regions under the conditions of DMSO and Nup62 depletion (ΔNup62); right panels show representative top views of assembled NPCs in each condition. Statistical significance applies to all panels in this figure: *, P ≤ 0.05; **, P ≤ 0.01; ***, P ≤ 0.001; ****, P ≤ 0.0001; ns, not significant.

Live imaging of cells expressing mScarlet-Nup58 revealed that Nup58, a direct binding partner of Nup62 in the Nup62-58-54 subcomplex, was absent from NPCs from the beginning of assembly when Nup62 was depleted (Fig. 2B; Fig. 2 Suppl. 2A), indicating that recruitment of the whole central channel subcomplex requires Nup62. Using immunofluorescence to detect Nups from different subcomplexes relative to Nup153 signal (Fig. 2 Suppl. 1B), we found that partially assembled NPCs in Nup62-depleted cells lacked its binding partners Nup214 and Nup88 in the Nup214-88-62 complex, which is involved in cytoplasmic export, as well as the nuclear basket protein Tpr (Fig. 2C-E). All these components were absent throughout the 5-30 min time window we sampled after AO (Fig. 2 Suppl. 2B), indicating that their recruitment is defective from the beginning of NPC assembly, rather than the loss of these proteins from initially assembled NPCs. In contrast, the recruitment of other key Nups, including the outer ring components Elys and Nup133, the inner ring Nup155, and the cytoplasmic filament Nup358, proceeded largely unperturbed. Only Nup98 showed a partial reduction (Fig. 2 Suppl. 3). These results indicate that Nup62 is specifically required for recruiting its direct binding partners in the central channel and on the cytoplasmic face, as well as the nuclear basket protein Tpr. In contrast, it is dispensable for the recruitment of major parts of the structural scaffold, including the outer and inner rings and cytoplasmic filaments.

We next asked whether the absence of specific Nups results from defective recruitment or destabilization of these proteins in the absence of Nup62. Quantitative western blotting of mitotically arrested cells revealed that the abundance of Nup214, Nup88, Nup58, and Nup54, all members of the Nup62 subcomplexes, was reduced to ∼60% of control levels while Nup62 itself dropped to ∼10% (Fig. 2 Suppl. 4A-C). By contrast, Nups outside these subcomplexes (Nup153, Nup358, Nup188, and Nup107) were not affected (Fig. 2 Suppl. 4D and E). In addition, depletion of Nup153 did not alter the protein levels of Nup62-binding partners (Fig. 2 Suppl. 4B-C). This partial reduction of the abundance of Nup62-binding partners is likely caused by the destabilization of the normally mitotically stable subcomplexes ^48–50^, upon Nup62 degradation. However, because more than half of these proteins remained stable, their failure to localize to the NE indicates Nup62 is required for their targeting to assembling NPCs.

Collectively, our findings demonstrate that Nup62 is specifically required for allowing the assembly of key molecular components of the NPC. Its acute depletion arrests postmitotic NPC assembly at an intermediate state (Fig. 2F) that lacks the central Nup62-58-54 channel subcomplex, the cytoplasmic Nup214-88-62 subcomplex, and the nuclear basket component Tpr. These incomplete pores also exhibit a significantly reduced diameter. Thus, Nup62 is essential for completing the molecular assembly and spatial dilation of the NPC after mitosis.

### Cohesive interactions between FG repeats in the Nup62-58-54 subcomplex explain its requirement for pore dilation during NPC assembly

The loss of the entire Nup62-58-54 subcomplex, which forms the central channel of the NPC at the neck of the nuclear membranes that failed to dilate, motivated us to explore how it might generate the necessary force to drive pore dilation (Fig. 3A). Structurally, this subcomplex is dominated by intrinsically disordered FG-repeat regions known to be essential for selective nuclear transport in the fully assembled NPC ^37–40^. For this transport function, multiple copies of this complex are proposed to interact with each other through multivalent, hydrophobic interactions between their disordered FG-repeat regions ^30–36,51^. However, whether these interactions also result in pore dilation during NPC assembly remains unknown.

**Figure 3.**
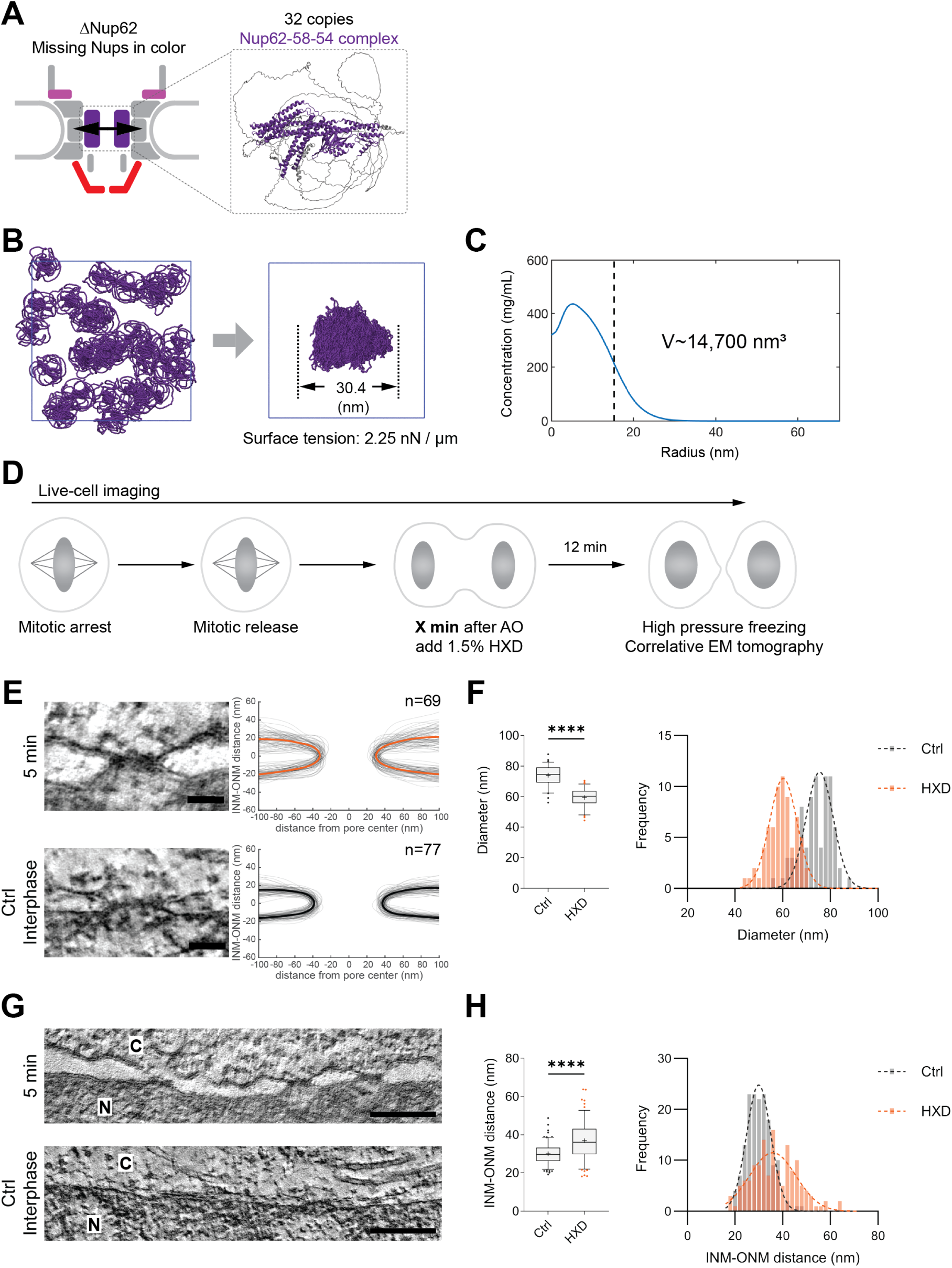
Acute disruption of FG-Nup interactions in the central channel during NPC assembly prevents pore dilation. **(A)** NPC schematic illustrating missing Nups (labeled in color) upon Nup62 depletion during mitosis. The AlphaFold-predicted heterotrimeric structure of the Nup62-58-54 subcomplex is shown, with the ordered domains in purple and the disordered regions in grey. **(B)** Molecular simulation with 32 copies of the Nup62-58-54 complex at 100 μM. Within 5 μs, all copies of the Nup62-58-54 subcomplex self-associated, resulting in an approximately spherical condensate with a diameter of 30.4 nm and a surface tension of 2.25 nN/µm. **(C)** Plot of the average simulated condensate concentration from (B) as a function of distance from the cluster center. The average concentration is taken from 12 independent simulations. The dashed line indicates the condensate radius, corresponding to a volume of ∼14,700 nm³. **(D)** Experimental scheme illustrating acute disruption of FG-Nup interactions during NPC assembly, correlating live imaging with electron tomography. Cells were treated with 1.5% 1,6-hexanediol (HXD) at defined time points (indicated by X min) after AO, incubated for 12 minutes, and subsequently high-pressure frozen for EM analysis. (**E)** Tomographic slices showing cross-section views of nuclear pores assembled in HK WT cells treated with HXD at 5 min after AO, as described in (D). NPCs from untreated interphase cells are shown for comparison. Membrane profiles of all measured pores are displayed, with mean profiles highlighted in bold. Scale bars, 50 nm. **(F)** Quantification of the NPC diameter based on (E). Left, box-and-whisker plot showing median, mean (“+”), 5–95 percentiles, and outliers (scatter). Right, histogram with Gaussian fits (dashed curves). Sample sizes as indicated in (E). **Note**: Minor differences in NPC diameter compared to Fig. 1 reflect slight variability in shrinkage during freeze substitution. Control samples were always processed in parallel with HXD-treated cells in each preparation batch. (**G)** Representative tomographic slices of the NE from the same samples as in (E). NE from untreated interphase cells is shown for comparison. N, nucleus; C, cytoplasm. Scale bars, 200 nm. **(H)** Quantification of the INM-ONM distance from cells in (G). Left, box-and-whisker plot showing median, mean (“+”), 5–95 percentiles, and outliers (scatter). Right, histogram with Gaussian fits (dashed curves). Sample sizes as indicated in (E).

To investigate if the hydrophobic interactions between FG-repeats in multiple copies of the Nup62-58-54 complex could generate a membrane bending force, we performed coarse-grained molecular dynamics simulations. These simulations focus on the collective properties of individual FG-Nups in solution, a condition under which some FG-Nups have been known to undergo phase separation experimentally ^30–37^. Using a modified version of the maximum entropy optimized force field (MOFF) ^52^, we quantified the phase separation propensity for the hydrophobic FG-repeat regions of Nup62 and its interacting partners, Nup58 and Nup54 (Fig. 3 Suppl. 1A-C). Indeed, the FG-repeat regions of Nup62, Nup58, and Nup54 were able to drive phase separation. In contrast, those of Nup214, which also fails to be recruited after Nup62 depletion, were not able to drive phase separation (Fig. 3 Suppl. 1C). Additional simulations confirmed that the full-length Nup62-58-54 subcomplex also had a clear phase-separation propensity (Fig. 3 Suppl. 1D).

To study the impact of Nup62-58-54 recruitment on an individual assembling nuclear pore, we simulated “condensates” formed by 32 copies of the Nup62-58-54 subcomplex, reflecting their stoichiometry within a single mature NPC ^19,21,22^. While it is unclear if such a condensate actually exists during the NPC assembly process, the simulations may be informative about the potential impact of cohesive FG-Nup interactions on the nuclear envelope during NPC assembly. At 100 µM concentration, the Nup62-58-54 subcomplex readily formed a globular condensate within 5 µs. The condensate had a surface tension of 2.25 nN/µm (Fig. 3B), which is well above theoretical estimates of the forces required to bend intracellular membranes ^53,54^. Further, the predicted volume of the condensate composed of 32 copies of the Nup62-58-54 subcomplex (∼14,700 nm³, Fig. 3B and C) was similar to the volume deficit we had observed by electron tomography in Nup62-depleted pores (∼15,400 nm³, Fig. 1B and C). Thus, our simulations predict that central channel Nups, when recruited into the previously unoccupied central region of the pore, could self-assemble. It is conceivable that the formation of such an assembly would exert an outward force to its surroundings, and that this force would be sufficient to drive pore dilation during NPC maturation.

To test this hypothesis in vivo, we acutely disrupted FG-FG hydrophobic interactions using 1,6-hexanediol (HXD), a reagent commonly used to disrupt FG-Nup interactions ^33,37,55^. For use in living cells, we established a minimal dose that disrupts FG-Nup function without broadly compromising cell viability. After careful titration, we found that 1.5% HXD rapidly compromised nuclear import in interphase cells, but had little effect on overall nuclear morphology (Fig. 3 Suppl. 2).

We next applied 1.5% of HXD to cells at 5 min after AO to acutely disrupt hydrophobic interactions just before pore dilation occurs^3,10^. We then, incubated cells for 12 min, the time required to complete NPC assembly ^3,4,10^, before fixing them for high-resolution imaging (Fig. 3D). Correlative ET revealed that pores formed under these conditions were ∼20% narrower than mature interphase pores (Fig. 3E and F; Fig. 3 Suppl. 3A), closely resembling the phenotype observed after Nup62 depletion (Fig. 1B and C; Fig. 1 Suppl. 2C and E; Fig. 1 Suppl. 3A). Similar results were obtained when HXD was added at 6 or 8 min after AO, just before or during pore dilation, while HXD additions no longer had an effect on pore diameter if added later than 13 min after AO, when dilation is normally complete ^3,4,10^, or during interphase (Fig. 3 Suppl. 3B and D). This data shows that hydrophobic interactions are essential for full pore dilation precisely during the time window when dilation occurs during unperturbed NPC assembly, but are no longer needed to maintain a normal NPC diameter once dilation is completed.

To rule out that HXD simply prevents recruitment of Nup62 to assembling NPCs rather than perturbing its hydrophobic interactions with other FG-Nups, we imaged its incorporation kinetics into the NE after mitosis. This kinetics was comparable to that in control cells, showing only a slight reduction in the final concentration reached at later times after AO (Fig. 3 Suppl. 4A and B). Thus, inhibition of pore dilation most likely results specifically from disruption of hydrophobic interactions rather than failure to target Nup62 to the NE after mitosis. Furthermore, HXD-treated daughter nuclei showed impaired nuclear import and were smaller in size than untreated nuclei (Fig. 3 Suppl. 4A, C and D), consistent with effective disruption of the selective nuclear transport barrier formed by hydrophobic FG-Nups after mitotic exit.

Together, our simulations and acute perturbation experiments support the model that hydrophobic FG-repeat interactions of the Nup62-58-54 subcomplex play an essential role in driving membrane pore dilation, a critical step in postmitotic NPC maturation.

### Import-induced nuclear expansion drives regular nuclear membrane spacing during mitotic exit

A striking correlation emerged from both Nup62-depleted and HXD-treated cells: their nuclear pores failed to dilate, while their NE also exhibited an increased and less regular spacing between the inner and outer nuclear membranes (Fig. 1D and E; Fig. 1 Suppl. 2D and F; Fig. 1 Suppl. 3B; Fig. 3G and H; Fig. 3 Suppl. 3C and E). During normal mitotic exit, the NE transitions from an initially loosely spaced double membrane derived from ER sheets to a parallel-aligned, tightly spaced structure right after NPC maturation ^3^. This temporal concurrence, as well as the perturbation of both pore dilation and nuclear membrane spacing in Nup62-depleted and HXD-treated cells, suggested that both processes might be mechanistically linked.

To explore this hypothesis, we looked at other features of the specifically perturbed cells. Notably, cells lacking Nup62 exhibited significantly smaller daughter nuclei (Fig. 4A), primarily due to inefficient nuclear import, as evidenced by the impaired nuclear accumulation of DiHcRed-NLS and the chromatin modulator Rad21 (Fig. 4 Suppl. 1A-C). Similarly, HXD-treated cells also led to import defects and smaller nuclei, despite normal incorporation of central FG-Nups (Fig. 3 Suppl. 4). This observation led us to hypothesize that nuclear import-mediated accumulation of nuclear proteins promotes nuclear expansion, which would generate tension in the nuclear membranes that promotes their progressively parallel alignment and tighter spacing after mitotic exit.

**Figure 4.**
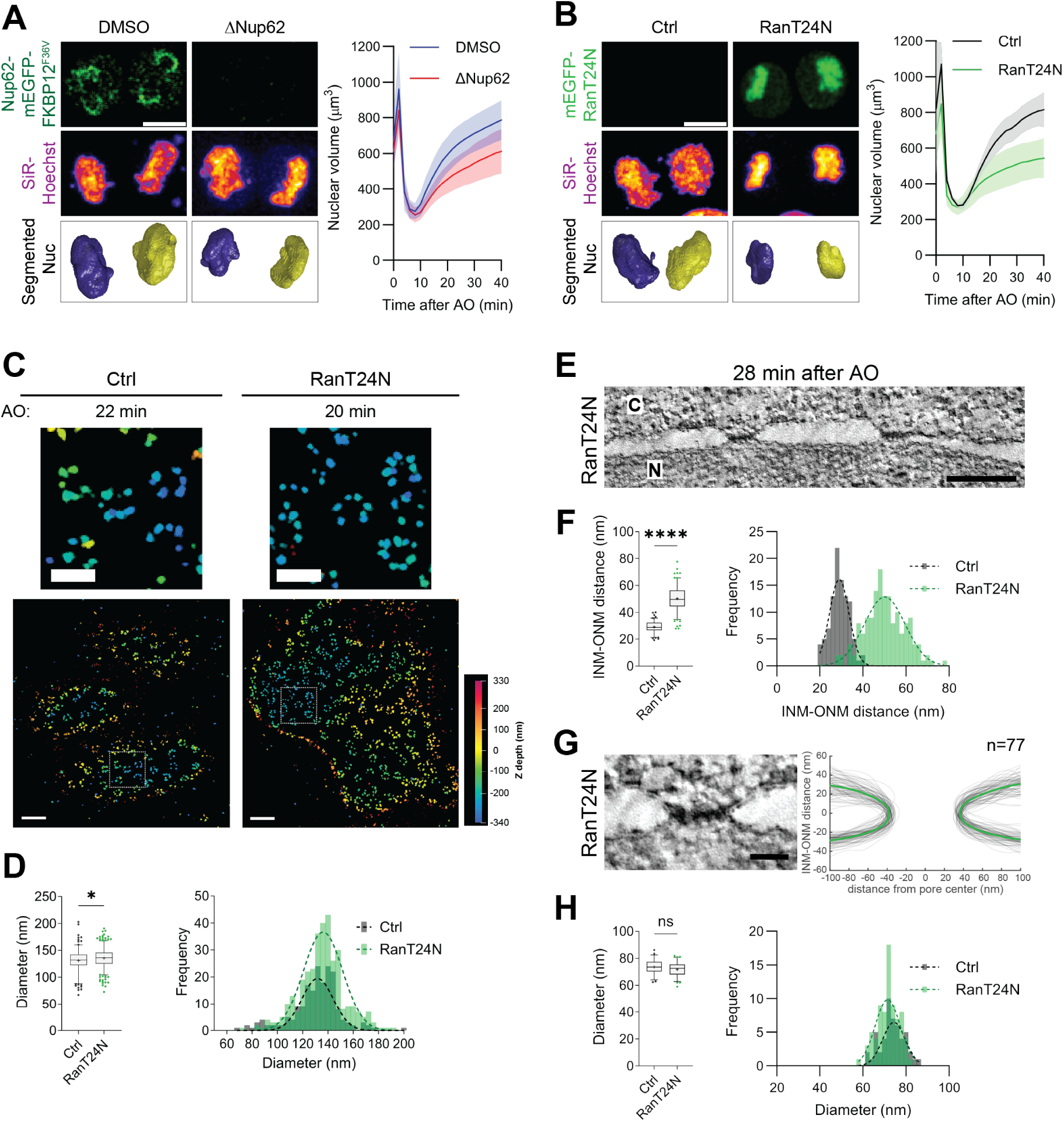
Acute import inhibition during mitotic exit increases NE spacing without affecting NPC diameter. **(A)** Representative live-cell images of HK Nup62-mEGFP-FKBP^F36V^ cells treated with DMSO or dTAG (ΔNup62), shown at 20 min after AO. Top, single z-slice of Nup62-mEGFP signal; middle, maximum projection of SiR-Hoechst-stained nuclei; bottom, 3D-segmented nuclei. Scale bar, 10 μm. Right, kinetic plot showing average nuclear volume during mitotic exit under the indicated conditions. Error bands represent the standard deviation. Sample size: DMSO: n = 41; ΔNup62: n = 22. **(B)** Representative live-cell images of HK WT cells under acute import inhibition (RanT24N), shown at 20 min after AO. The workflow is described in Fig. 4 Suppl. 1B. Ctrl cells lacked mito-LAMA and mEGFP-RanT24N expression but were treated with TMP. Top, single z-slice of mEGFP-RanT24N signal after release; middle, maximum projection of SiR-Hoechst-stained nuclei; bottom, 3D-segmented nuclei. Scale bar, 10 μm. Right, kinetic plot showing average nuclear volume during mitotic exit under the indicated conditions. Error bands represent the standard deviation. Sample size: Ctrl, n=7; RanT24N, n=21. **(C)** Representative 3D MINFLUX images (anti-Elys) of cells under Ctrl or RanT24N conditions at the indicated time points after AO. Scale bars: upper panel, 200 nm; lower panel, 500 nm. **(D)** Quantification of the NPC diameter based on (C), from cells dividing between 20 and 60 min after AO. Left, box-and-whisker plot showing median, mean (“+”), 5–95 percentiles, and outliers (scatter). Right, histogram with Gaussian fits (dashed curves). Sample sizes: Ctrl, n=191; RanT24N, n=340. **(E)** Representative tomographic slice of the NE in HK WT cells under RanT24N condition at 28 min after AO. N, nucleus; C, cytoplasm. Scale bar: 200 nm. **(F)** Quantification of the INM-ONM distance from cells in (E), with interphase Ctrl cells included for comparison. Left, box-and-whisker plot showing median, mean (“+”), 5–95 percentiles, and outliers (scatter). Right, histogram with Gaussian fits (dashed curves). (**G)** Left, tomographic slices showing cross-section views of nuclear pores from the same samples as in (E). Right, membrane profiles of all measured pores are displayed, with mean profiles highlighted in bold. Scale bar, 50 nm. **(H)** Quantification of the NPC diameter based on (G), including interphase Ctrl cells for comparison. Left, box-and-whisker plot showing median, mean (“+”), 5–95 percentiles, and outliers (scatter). Right, histogram with Gaussian fits (dashed curves). Sample sizes as indicated in (G) and Fig. 4 Suppl. 2D. Statistical significance applies to all panels in this figure: *, P ≤ 0.05; **, P ≤ 0.01; ***, P ≤ 0.001; ****, P ≤ 0.0001; ns, not significant.

To test this idea, we devised a system to acutely inhibit nuclear import in living cells during mitotic exit (Fig. 4 Suppl. 1D). Adapting a chemogenetic ligand-modulated antibody fragment (LAMA) system ^56^, we sequestered a dominant-negative mutant of the central regulator of nucleocytoplasmic transport Ran (RanT24N) away from the nucleus at the mitochondria, allowing cells to enter mitosis normally. Using the specific ligand, trimethoprim (TMP), to release RanT24N bound to the LAMA system, we could then trigger the rapid targeting of the dominant-negative mutant to mitotic chromosomes just before mitotic exit (Fig. 4 Suppl. 1D). As expected, mitotic RanT24N release acutely blocked nuclear import and arrested nuclear expansion during mitotic exit (Fig. 4B; Fig. 4 Suppl. 1E-G), closely resembling the phenotypes observed in Nup62-depleted and HXD-treated cells (Fig. 4A; Fig. 4 Suppl. 1A-C).

We next studied whether the RanT24N-mediated block in nuclear import might also have perturbed NPC assembly. Correlative 3D MINFLUX imaging of Elys-labeled pores revealed no dilation defects of the outer ring of the nuclear pore after RanT24N release up to one hour after AO (Fig. 4C and D; Fig. 4 Suppl. 2A). Thus, acute nuclear import inhibition does not appear to impair NPC assembly, but specifically blocks transport activity and thereby nuclear expansion. To test if nuclear membrane spacing is affected by a lack of nuclear expansion, as we had hypothesized, we then examined the nuclear membrane topology ultrastructurally using correlative ET. Interestingly, RanT24N-inhibited cells indeed exhibited significantly increased INM-ONM spacing by 28 min after AO (Fig. 4E and F; Fig. 4 Suppl. 2B and C). In contrast, control cells observed at a similar time point had already adopted a tightly parallel, interphase-like NE architecture (Fig. 1 Suppl. 2D and F). Importantly, measuring the diameter of the membrane pore necks in individual assembled NPCs confirmed that pores in RanT24N-inhibited cells exhibited diameters comparable in size to those in control cells (Fig. 4G and H; Fig. 4 Suppl. 2D), confirming our 3D MINFLUX result that pore dilation is not affected by blocking nuclear transport (Fig. 4C and D; Fig. 4 Suppl. 2A).

Taken together, these observations demonstrate that Ran-mediated nuclear import is not required for NPC dilation. However, Ran-mediated import is required for nuclear expansion and establishing a tight and regular parallel spacing between the two nuclear membranes. The coupling between nuclear pore dilation and nuclear membrane spacing can therefore be dissected as a two-step mechanism, as failure in condensate formation and nuclear pore dilation does not allow the assembly of import-competent pores, which in turn does not allow nuclei to expand and generate the tension required to tighten their double membranes.

## Discussion

The molecular logic linking postmitotic NPC assembly and NE remodeling has remained elusive. Here, we show how the two processes can be linked *via* an interplay between multimolecular self-assembly and biophysical force generation. We identify the central channel FG-Nups as active agents of pore dilation during nuclear boundary formation after mitosis. Upon self-assembly, their multivalent, hydrophobic interactions enable FG-Nups to populate the previously unoccupied central region of the pore, which transforms the small membrane opening in the ER-derived sheets on the chromosome surface into a transport-competent and structurally stable nuclear pore. This step, in turn, enables import-driven nuclear expansion, which generates the mechanical force across the whole nuclear surface to reshape the NE into a flat surface with a tightly spaced parallel double membrane boundary after mitosis (Fig. 5).

**Figure 5.**
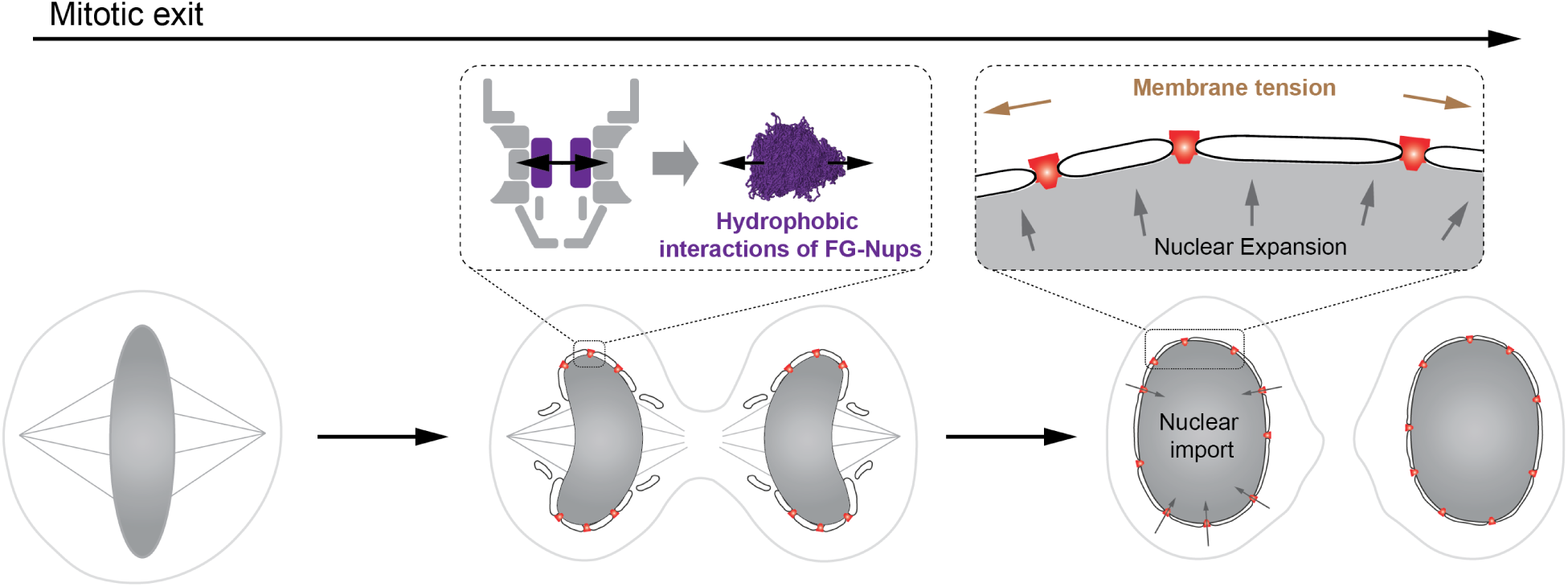
A unifying model of NPC assembly and NE remodeling during mitotic exit. See discussion.

### Central FG-Nups are essential for pore dilation during NPC maturation

Our temporally resolved compositional and structural analysis revealed that acute Nup62 depletion during mitosis halts NPC assembly at an early stage with a narrowed membrane pore diameter, which lacks key molecular components, including the central transport channel Nup62-58-54 subcomplex, Nup214-88-62 subcomplex, and the nuclear basket component Tpr (Fig. 1 and 2). This smaller pore is in contrast to the widened pores that have been seen upon depletion of outer ring Nups (e.g., Nup96 or Nup133) in interphase ^54,55^. These observations demonstrate a central role for Nup62 and its binding partners in mediating pore dilation.

Correlative immunofluorescence revealed that Nup62 is dispensable for the initiation of NPC assembly by the earliest recruited Nups, including Nup153. Instead, Nup62 acts right after initiation, as a critical assembly hub for the compositional maturation of the pore. The failure to recruit the Nup62-58-54 central channel and Nup214-88-62 cytoplasmic subcomplex indicates that Nup62 is central to orchestrating the assembly of its binding partners ^56,57^ in these subcomplexes into the assembling NPC, as they only showed a minor degree of co-depletion ^52,53^. On the other hand, impaired Tpr recruitment, which is not reported to interact directly with Nup62, is most likely due to the inability to establish its nuclear import in the absence of Nup62 (Fig. 4 Suppl. 1A-C) ^57,58^. Consistent with this hypothesis, recruitment of the Nup358 cytoplasmic filament remained unaffected, despite its late incorporation ^4^, as it does not require nuclear transport but only anchoring to the cytoplasmic outer ring complexes ^20,59^.

Structural analysis of the assembly intermediates by correlative ET further illuminated the essential role of Nup62 and its associated Nups for the structural maturation of the pore. The arrested, constricted NPCs showed a ∼21% reduction in diameter of their membrane neck, corresponding to an almost 40% reduction of the cross-section available for transport. Interestingly, Elys-based MINFLUX measurements of the outer nuclear ring showed a ∼5% decrease, revealing differences in flexibility of different parts of the NPC: while the inner ring is malleable, the outer ring appears relatively rigid, in line with previous findings on mature NPC structural flexibility ^46,47^. These findings support a model in which the outer nuclear ring forms a stabilizing scaffold early during postmitotic NPC assembly, in turn providing the stable template for subsequent formation of the Nup62 subcomplexes that then mediate dilation and maturation of the inner pore. These results validate our previous integrative spatiotemporal model of NPC assembly ^4,10^.

### Hydrophobic interactions between central channel FG-Nups promote pore dilation

Mechanistically, we propose that the recruitment of the three central channel FG-Nups together with the hydrophobic, multivalent interactions between their FG repeats drives pore dilation by generating the outward pushing force. This hypothesis is supported by molecular dynamics simulations that predict condensation of the central channel FG-Nups Nup62-58-54 in solution (but importantly not of the cytoplasmic FG-Nup214) at physiologically relevant stoichiometry and concentration. Comparison with the yeast and plant orthologs of the central channel subcomplex shows that the biophysical properties of these FG-Nups are evolutionarily conserved ^31,37,51^. The simulations further predict that the capillary pressure generated by such small condensates, determined by surface tension and their radius, can exert an outward pushing mechanical force to its surroundings ^60^. Although it is unclear if such a condensate exists in the actual assembling NPC, the early presence of the Nup62-58-54 complex indicates that similar hydrophobic, multivalent FG-Nup interactions are present throughout postmitotic assembly, and their establishment in the previously unoccupied central region of the pore could generate the force necessary for pore dilation. Indeed, in other membrane deformation contexts, interactions between disordered proteins generate the forces necessary for membrane budding, tubulation, and even scission ^53,54,61–65^. In summary, our simulations predict that FG-Nup interactions within the Nup62-58-54 subcomplex would generate sufficient force to physically push the neck of small nuclear membrane openings apart, thus dilating the nuclear pore during its assembly. It is also conceivable that this Nup62-58-54 subcomplex is targeted to the small nuclear membrane pore by the previously assembled structurally stable nuclear outer ring.

We experimentally tested the computational predictions in vivo by disrupting hydrophobic interactions during NPC assembly with HXD. At a minimal dose, HXD specifically disrupts hydrophobic FG-Nup interactions, while not impairing Nup62 recruitment, consistent with prior yeast studies showing that HXD has minimal effects on Nup localization, stability, and aggregation ^55,66^. Strikingly, HXD, if applied precisely during the time window of NPC maturation, recapitulated the pore dilation defect seen in Nup62-depleted cells. Importantly, if HXD was applied a few minutes after the completion of NPC structural maturation, it no longer had an effect (Fig. 3 and Fig. 3 Suppl. 2). This assembly stage-specific sensitivity of pore dilation to HXD treatment in live cells provides strong support for the hypothesis that hydrophobic interactions of the central channel FG-Nups are driving pore dilation.

### Import-driven nuclear expansion flattens the double membrane boundary of the nucleus

The morphological transition from loosely spaced postmitotic nuclear membranes derived from ER sheets to the tightly parallel-aligned flat interphase architecture of the nuclear surface occurs shortly after postmitotic NPC assembly ^3^. In Nup62-depleted or HXD-treated cells, this transition was impaired, and nuclear membrane spacing remained abnormally wide and irregular (Fig. 1 and 3). Because HXD-treated cells still recruited Nup62 at NPCs, these findings suggest that it is not the mere presence of Nup62, but the hydrophobic interactions between the FG repeats, that are required for NE remodeling.

We also addressed the question of why the generated forces that locally dilate individual membrane pores also flatten the whole NE. Our finding that both Nup62 depletion and HXD treatment impaired nuclear import and led to smaller nuclei (Fig. 4 and Fig. 4 Suppl. 1) provided the key for an explanation, which is that it is not pore dilation, but rather the resulting core function of the assembled NPC, nuclear import, that is needed for flattening the NE. Our system to acutely activate RanT24N during mitosis allowed us to dissect these two Nup62-dependent steps. This specific and acute perturbation showed that inhibition of nuclear import, without affecting NPC assembly, is sufficient to generate smaller daughter nuclei and cause increased and irregular nuclear membrane spacing. These findings collectively support a broader principle: postmitotic nuclear import drives nuclear expansion, which in turn remodels the NE. This principle is consistent with previous findings in cells with defects in other NPC components or Ran pathway regulators ^14,15,41,67^.

It has been proposed previously that nuclear expansion might generate the entropic outward-pushing mechanical forces stretching the nuclear membrane ^46,47,68^. Computational models estimated that ∼0.15 nN/µm of tension is sufficient to maintain a flat NE architecture, while reduced tension results in increased spacing ^69^. Our data from all perturbation conditions (Nup62 depletion, HXD treatment, and acute import inhibition during mitotic exit) are consistent with this model, reinforcing the link between nuclear import, nuclear expansion, and NE remodeling. Further supporting this link, increased membrane tension has been correlated with reduced NE spacing during neuronal differentiation ^70^.

Membrane tension has also been proposed to regulate the diameter of fully assembled NPCs in interphase nuclei. It has indeed been shown that increased tension can mechanically stretch the NPC scaffold to dilate the pore beyond its normal diameter, whereas reduced tension results in a constriction ^46,47,70,71^. For NPC assembly after mitosis, however, which occurs into still unstretched irregular nuclear membranes derived from the ER and before the onset of nuclear import, our data clearly shows that pore dilation requires Nup62 and hydrophobic interactions, but not nuclear import, because preventing the buildup of membrane tension by import inhibition does not impair pore dilation during NPC assembly (Fig. 4). Molecular dynamics simulations furthermore suggest that the surface tension of individual FG-Nup condensates (∼2.25 nN/µm) is over an order of magnitude larger than the estimated nuclear membrane tension (∼0.15 nN/µm) ^69^, and thus could provide the driving force for pore dilation during postmitotic assembly.

### A unifying model of NPC assembly and NE remodeling

Together, our results allow us to propose a novel unifying mechanistic model of NPC assembly and nuclear membrane remodeling after mitosis. In this model, FG-Nup cohesive interactions, particularly between the FG repeats in the Nup62-58-54 complex, generate localized forces that drive pore dilation. This structural transition enlarges the membrane channel and establishes the transport barrier, thereby enabling nuclear import, which in turn drives nuclear expansion, increasing nuclear membrane tension and generating the mechanical forces to shape the NE into its flat interphase architecture with tightly spaced parallel nuclear membranes. Distinct phenotypes, such as blocked pore dilation, impaired import, reduced nuclear expansion, and nuclear membrane disorganization, result from how this sequence of events is disrupted (by depletion of Nup62 prior to dilation, disruption of FG-Nup interactions during dilation, or inhibition of import after pore dilation and formation of the transport barrier (Fig. 5). Our work thus provides a mechanistic understanding of how mechanical forces first generated locally by hydrophobic FG-Nup interactions and then across the whole nuclear surface by nuclear import are used in a temporally coordinated fashion during nuclear reformation after mitosis.

It is tempting to speculate that a related mechanism may also be used to promote the assembly of new NPCs into the interphase nucleus. Here, FG-Nup interactions might promote the local evagination of the inner nuclear membrane that has been observed prior to the de novo fusion needed to insert NPCs into the closed double membrane ^1,72^. More broadly speaking, our findings suggest that nanoscopic assemblies of 10s-100s copies of hydrophobic, disordered proteins can act as local force generators to plug and dilate existing small membrane pores, or to bend them outwards to bring them into sufficiently close proximity for fusion. This model provides a simple mechanistic principle, based on local self-assembly, for how intrinsically disordered hydrophobic regions may have contributed to the emergence of intracellular compartmentalization, hinting at an ancestral role for hydrophobic interaction-driven mechanics in the evolution of the eukaryotic nucleus.

## Acknowledgments

We thank the Electron Microscopy Core Facility for their support in correlative light and electron microscopy (especially R. Mellwig, V. Oorschot, and M. Schorb), and the members of the Ellenberg group (especially A. Brunner) and Shotaro Otsuka for advice and discussion. We acknowledge the access and services provided by the Imaging Centre at the European Molecular Biology Laboratory, generously supported by the Boehringer Ingelheim Foundation. We gratefully acknowledge Merle Hantsche-Grininger for reading and providing feedback on the manuscript. The work was supported by grants from the Baden-Württemberg Foundation (J.E. and A.S.); the Deutsche Forschungsgemeinschaft (DFG, German Research Foundation) -Project-ID 511488495 -SFB 1638, P06 to J.E.; NIH/NIGMS R01GM083960 (A.S.), NIH/NIGMS P41GM109824 (A.S.), NIH/NIGMS R01GM112108 (A.S.); the European Molecular Biology Laboratory (W. Z., N.R.M., H.P., M.H.-G., and J.E.). A.P.L. was further supported by NIH/NIGMS F32GM150243. This work used Expanse GPUs at the San Diego Super Computer through allocation BIO230055 from the Advanced Cyberinfrastructure Coordination Ecosystem: Services & Support (ACCESS) program, which is supported by U.S. National Science Foundation grants #2138259, #2138286, #2138307, #2137603, and #2138296.

## Author contributions

W.Z. and J.E. conceived and designed the project. W.Z. performed the live-cell imaging, immunofluorescence, and Simple Western experiments with the assistance from N.R.M. and H.P. A.P.L. contributed to the computational simulation of condensate formation. W.Z. performed correlative ET together with P.R., and correlative MINFLUX imaging together with S.S.. Most data analyses were performed by W.Z.; 3D MINFLUX data analysis was conducted by J.K.H.. Z.H. contributed to the computational quantitative analysis of EM images and MINFLUX data preprocessing. M.J.H. contributed to the segmentation of fluorescence images. Y.S., A.S., and J.E. supervised the work. W.Z. and J.E. wrote the paper. M.H.-G. and N.R.M. revised the initial draft of the manuscript. W.Z. and J.E. revised and edited the manuscript with feedback from all authors. All authors discussed the results and approved the final version of the manuscript.

## Methods

### Cell culture

HeLa Kyoto wild type (HK WT, S. Narumiya (Kyoto University, Kyoto, Japan), RRID: CVCL_1922) and all derivative HK cell lines were cultured in high-glucose DMEM (Thermo Fisher Scientific, 41965-062) supplemented with 10% FBS (Thermo Fisher Scientific, 10270-106), 100 U/ml penicillin-streptomycin (Thermo Fisher Scientific, 15140-122) and 1 mM sodium pyruvate (Thermo Fisher Scientific, 11360-039) at 37°C, 5% CO_2_ unless otherwise stated. Cells were grown in cell culture dishes or flasks and passaged every 2-3 days via trypsinization with 0.05% Trypsin-EDTA (Thermo Fisher Scientific, 25300-054). Mycoplasma contamination was checked regularly using a PCR-based detection kit (abm, G238) and confirmed to be negative.

### Plasmid transfection

Plasmids constructed and used in this study are listed in the Supplementary Table 1. Transient transfections were performed using polyethylenimine (PEI) transfection reagent (1 mg/ml stock, Polysciences, 24765-1), and cells were incubated for 48 - 72 h before imaging. For 8-well chambered slides, 0.1μg of plasmid and 0.4μL of transfection reagent in 20 μL of opti-MEM Serum-Reduced Medium (Thermo Fisher Scientific, 31985070) were used for transfection per well. For 35 mm gridded dishes, 1 μg of plasmid and 4 μL of transfection reagent in 200 μL of opti-MEM Serum-Reduced Medium were used.

### Generation of homozygous endogenous knock-in cell line

The genome-edited cell line generated for this study (HK Nup62-mEGFP-FKBP12^F36V^ #C02) was created by C-terminal tagging of Nup62 in the HK WT parental cell line using the CRISPR/Cas9 method. The detailed protocol was published previously ^42^. In brief, a linear DNA donor sequence encoding for the tag of interest, including 40 bp homology arms, was electroporated (Neon Transfection System, Thermo Fisher Scientific) into the parental HK cell line, together with the catalytic Cas9/gRNA ribonucleoparticle complex. For this, we used Alt-R S.p. HiFi Cas9 Nuclease V3 (IDT, 1081061) and single gRNAs (see Supplementary Table 2).

### Cell synchronization to prometaphase

To synchronize cells in prometaphase for subsequent protein degradation and/or live imaging, two different approaches were used. For brief synchronization, cells were treated with either 82.5 nM nocodazole (5 mg/ml stock, Sigma, SML1665) or 2.5 μM STLC (50 mM stock, Sigma, 164739) for 3-4 h. For strong enrichment of the prometaphase population, cells were first arrested with 2 mM thymidine (Sigma, T-1895) for 24 h, followed by 12–14 h of STLC (2.5 μM) treatment.

### Acute protein depletion during mitosis

Genome-edited HK cells expressing endogenously tagged Nup62 (Nup62-mEGFP-FKBP12^F36V^) and Nup153 (Nup153-mEGFP-FKBP12^F36V^) ^14^ using the dTAG degron system were used for the degradation of Nup62 and Nup153, respectively. The depletion of degron-tagged proteins was induced in the presence of 82.5 nM nocodazole or 2.5 μM STLC, along with degradation-triggering compounds (dTAG-13 [Sigma-Aldrich, SML2601] and/or dTAG^V^-1 [Tocris, 6914]). To optimize depletion efficiency, a serial titration of the concentrations of dTAG compounds was performed. For all functional assays, cells were treated with 250 nM dTAG-13 and 500 nM dTAG^V^-1 for 90 min in the prometaphase-arrested stage. Following treatment, cells were released into mitosis by washing out nocodazole or STLC and replenishing with fresh medium containing dTAG compounds.

### Acute import inhibition assay

HK WT cells transiently co-expressing mito-LAMA and mEGFP-RanT24N were synchronized to prometaphase and subjected to live-cell imaging, following the procedure described in the “Live-cell imaging” section. The assay was conducted at 37 °C. Prometaphase-arrested cells were released into mitosis, and z-stack images were captured from multiple fields of view containing several cells at different positions. Subsequently, trimethoprim (TMP) (Sigma Aldrich, T7883) was added to a final concentration of 50 µM to induce the release of mitochondrial-tethered mEGFP-RanT24N into the nucleus. Timelapse imaging was recorded immediately after TMP addition to monitor mitotic progression.

### 1,6-hexanediol assay

For 1,6-hexanediol (HXD, Sigma Aldrich, 240117) treatment of interphase cells, Nup62-mEGFP-FKBP12^F36V^ cells transiently expressing DiHcRed-NLS were subjected to live-cell imaging as described in the “Live-cell imaging” section. Experiments were conducted at 37 °C. Before HXD treatment, z-stack images were acquired from multiple fields of view containing interphase cells. Subsequently, HXD was added directly to the imaging medium at a 1:1 volume ratio to achieve final concentrations of 1%, 1.5%, 2%, 3%, and 4% (w/v). Timelapse imaging was recorded immediately after HXD addition.

For HXD treatment of mitotic cells, HK WT or Nup62-mEGFP-FKBP12^F36V^ cells were synchronized at prometaphase and subjected to live-cell imaging. Before HXD treatment, cells were released into mitosis, and z-stack images were acquired from multiple fields of view containing cells undergoing mitotic exit. Subsequently, HXD was added to the imaging medium at a 1:1 volume ratio to achieve a final concentration of 1.5% (w/v). Timelapse imaging was recorded immediately after HXD addition to monitor mitotic progression.

### Live-cell imaging

Cells were seeded at least one day before live imaging on 8-well chambered cover glasses (ibidi µ-slide, 80807) for high-resolution live imaging or on 35 mm gridded dishes (ibidi µ-Dish, 81168) for correlative live imaging with immunofluorescence or 3D MINFLUX imaging. Both cover glasses and dishes were pre-coated with 0.1 mg/ml poly-L-lysine (Sigma-Aldrich, P-8920). One hour before live-cell imaging, DMEM medium was exchanged to phenol-red free CO_2_-independent imaging medium based on Minimum Essential Medium (Sigma-Aldrich, M3024) containing 30 mM HEPES (pH 7.4), 10% FBS, and 1x MEM non-essential amino-acids (Thermo Fisher Scientific, 11140-050). DNA dyes (50 - 100 nM of 5-SiR-Hoechst [gift from G. Lukinavičius ^73^] or Abberior LIVE 610 DNA [Abberior, LV610-0143]) were added to monitor mitosis.

Imaging was conducted at 37°C within a microscope-body-enclosing incubator. High-resolution 3D time-lapse imaging was performed using a confocal microscope (LSM780 or LSM880; Carl Zeiss) equipped with a C-Apochromat 40×/1.2 W Korr UV-Vis-IR water-immersion objective (Carl Zeiss). The cell division process was monitored every 1 or 2 min with 21-25 z-slices and a voxel size of 0.25 µm in xy and 1 µm in z, section thickness of 2.0 μm, covering a total of 61 × 61 µm in xy (240 × 240 pixels) and 20 µm in z. Fluorescence images were filtered with a mean filter (kernel size: 1 pixel) in ImageJ for presentation purposes. For correlative live imaging, cells were observed at a widefield microscope or confocal microscope (Axio Observer Z1 or LSM780; Carl Zeiss) using 20 × 0.4 NA Plan-Neofluar objective (Carl Zeiss) or 20 × 0.8 NA Plan-Apochromat objective (Carl Zeiss). The cell division process was monitored every 1 min.

### Correlative high-resolution immunofluorescence

Cells seeded on the 35 mm gridded dishes were monitored by live imaging to follow mitotic exit and then fixed with 2.4% paraformaldehyde (PFA) (EMS, 15710) in PBS for 15 min. PFA was quenched for 5 min with 100 mM NH_4_Cl in PBS. Permeabilization was performed with 0.25% Triton X-100 (Sigma Aldrich, T-8787) in PBS for 15 min. For immunolabeling, cells were incubated in blocking buffer (2% bovine serum albumin (Sigma Aldrich, A2153) and 0.05% Triton-X100 in PBS) for 30 min at room temperature (RT), followed by incubation with primary antibodies (listed in the Supplementary Table 3) diluted in blocking buffer at 4 °C overnight. After washing (3x for 5 min) in blocking buffer, cells were incubated with the secondary antibodies (listed in the Supplementary Table 3) diluted in blocking buffer for 30 min at RT. After multiple washes in PBS, cells were post-fixed with 2.4% PFA in PBS for 15 min and quenched for 5 min with 100 mM NH_4_Cl in PBS. Finally, the samples were washed 3 times for 5 min with PBS before the addition of 2 μg/ml Hoechst 33342 (Sigma Aldrich, B2261) in PBS to detect DNA. Cells of interest were identified from the recorded movies using the live DNA dye channel, and their positions were determined based on the grid pattern on the dishes using the T-PMT and Hoechst 33342 channels. Imaging was performed on a Zeiss LSM780 confocal microscope equipped with a C-Apochromat 40×/1.2 W Korr UV-Vis-IR water-immersion objective (Carl Zeiss). Images of fixed dividing cells were acquired in 35 z-slices with a voxel size of 0.07 µm in xy and 0.45 µm in z, covering a total of 35 × 35 µm in xy (512 × 512 pixels) and 15.3 µm in z. All immunofluorescence images shown in the study (unless otherwise stated) are from a single focal plane and were filtered with the PureDenoise plugin in ImageJ for presentation purposes.

### Light microscopy data analysis

#### Kinetic analysis in living cells

To assess degradation kinetics at varying concentrations of dTAG compounds, timelapse images acquired from single focused-plane Z-slices were analyzed in CellProfiler (version 4.2.5). A cell mask was generated from the 500 kDa dextran-Dy481XL fluorescence signal using CellProfiler built-in segmentation algorithms. This mask was then applied to quantify Nup62-mEGFP-FKBP12^F36V^ fluorescence intensity during depletion treatment. Mean fluorescence intensities were normalized to the baseline intensity measured immediately before dTAG compound addition.

To measure mScarlet-Nup58 mean fluorescence intensity during mitotic exit in the Nup62-mEGFP-FKBP12^F36V^ cell line treated with DMSO or dTAG compounds, maximum-intensity projections of 3D image stacks were analyzed using CellProfiler (version 4.2.5). NE and cytoplasmic masks were generated from mScarlet-Nup58 and DNA staining signals using CellProfiler built-in segmentation algorithms. These masks were then applied to quantify mScarlet-Nup58 fluorescence on the NE and in the cytoplasm. The ratio of mean NE-to-cytoplasmic fluorescence intensities was calculated to generate kinetic profiles during mitotic exit.

To evaluate the effects of different HXD concentrations on interphase cells, 3D Z-stack images were analyzed using a custom MATLAB script adapted from a previously published version ^4^. The script segmented the nucleus in 3D and quantified both nuclear volume and mean fluorescence intensity within the segmented region. These values were then normalized to baseline measurements acquired immediately before HXD addition.

To evaluate the effects of acute HXD treatment on postmitotic cells, 3D Z-stack images were analyzed using a previously published custom MATLAB script ^4^. The script segmented non-core regions of the postmitotic NE, from which mean fluorescence intensities of Nup62-mEGFP were extracted and baseline-corrected by subtracting the corresponding cytoplasmic mean intensity measured in the same cell during metaphase. These values were then normalized to the plateau mean intensities of Nup62-mEGFP in the segmented non-core regions of the postmitotic NE under control conditions after mitosis. In parallel, the script also segmented the entire nucleus in 3D to quantify nuclear volume, a procedure that was also applied for nuclear volume measurements under Nup62 depletion and RanT24N activation conditions.

To evaluate Nup62 depletion, acute HXD treatment or RanT24N activation on nuclear transport after mitosis, maximum-intensity projections of 3D image stacks from cells expressing DiHcRed-NLS were analyzed using CellProfiler (version 4.2.5). Nuclear and cytoplasmic masks were generated from DiHcRed-NLS and DNA staining signals using CellProfiler built-in segmentation algorithms. These masks were then applied to quantify DiHcRed-NLS fluorescence in the nucleus and cytoplasm. The ratio of mean nuclear-to-cytoplasmic fluorescence intensities was calculated to generate kinetic profiles during mitotic exit.

#### Quantification of correlative immunofluorescence

To quantify fluorescence intensities in the non-core regions of divided cells co-stained with anti-Nup153 and a second Nup antibody, 3D Z-stack images were analyzed using a custom MATLAB script adapted from a previously published version ^4^. The script segmented non-core regions of the postmitotic NE, from which mean fluorescence intensities of the second Nup antibody were extracted. These values were then normalized to the mean intensity of anti-Nup153 measured in the same segmented region.

To quantify the ratio of mean fluorescence intensities between the nucleus and cytoplasm in cells stained with anti-Rad21, maximum-intensity projections of 3D image stacks were analyzed using CellProfiler (version 4.2.5). Cytoplasmic and nuclear masks were generated from anti-Rad21 and DNA staining signals using CellProfiler built-in segmentation algorithms. These masks were then applied to quantify Rad21 fluorescence in the nucleus and cytoplasm. The ratio of mean nucleus-to-cytoplasmic fluorescence intensities was calculated.

### Correlative 3D MINFLUX DNA-PAINT imaging

Sample preparation followed the procedure described in the “Correlative high-resolution immunofluorescence” section. After blocking, cells were incubated with a primary antibody mixture of anti-ELYS antibody (polyclonal rabbit anti-AHCTF1 antibody; Sigma-Aldrich; HPA031658; 1:50) and Mab414 (Biolegend, 902902, 1:500) at 4 °C overnight, followed by incubation with a secondary nanobody mixture containing the anti-rabbit single-domain nanobody coupled to a DNA-PAINT sequence (Massive photonics, MASSIVE-sdAB 1-PLEX labeling kit, 1:100) and goat anti-mouse IgG, Alexa Fluor™ 555 (Invitrogen, A-21422, 1:5000) for 30 min at RT. Cells of interest were identified from the previously recorded movies, and their positions were determined based on the grid pattern on the dishes using the DIA channel via the Epi illumination on the MINFLUX microscope.

Gold nanoparticles (Cytodiagnostic, 150 nm, CG-150-100) were used as fiducials for sample stabilization during MINFLUX measurements ^74^. An undiluted dispersion of the nanoparticles was applied to the samples and incubated for 5 min at RT. The samples were then gently rinsed several times with PBS to remove unbound nanoparticles and then transferred to Imaging Buffer supplemented with 250 pM of DNA-PAINT imager strand F2-Atto655 (Massive photonics, MASSIVE-sdAB 1-PLEX labeling kit). 3D MINFLUX imaging was carried out using a commercial MINFLUX microscope (Abberior Instruments, Göttingen, Germany) equipped with a 100x oil immersion objective lens (UPL SAPO100XO/1.4, Olympus, Tokyo, Japan). We used the 642 nm CW excitation laser, two avalanche photodiodes (SPCM-AQRH-13, Excelitas Technologies, Mississauga, Canada) with a detection range of 650 - 685 nm for the first channel and 685 - 760 nm for the second (detected photons were summed), and a pinhole size corresponding to 0.69 airy units. All the hardware was controlled by Abberior Imspector software (v.16.3.15645-m2205-win64-MINFLUX, Abberior Instruments). Cells of interest identified from the Epi illumination were placed in focus using the Cy3 confocal reference channel. Before starting a MINFLUX measurement, the active sample stabilization system of the microscope was locked using gold nanoparticles deposited on the dish surface as positional references. Measurements were conducted with a stabilization precision of typically below 1 nm in xyz, respectively.

The MINFLUX acquisition is based on a MINFLUX sequence (set of parameters specified within a supplementary text file, see Imaging_3D.json), which controls the iterative zooming in on single-molecule events and was provided by the manufacturer. The MINFLUX iteration process is described in ^75^. In three dimensions, nine iterations plus one pre-localization iteration were performed. In the last iteration, a scan-pattern diameter (L-size) of 40 nm was used.

### MINFLUX data analysis

Each MINFLUX measurement was exported using Imspector Software (Abberior Instruments) as a MAT file. The exported files contained a collection of recorded parameters for all valid localizations. These files were then imported into a custom MATLAB software, MINFLUX data viewer (https://git.embl.de/grp-ic/minflux-data-viewer), for preprocessing. The preprocessing steps included: (1) scaling the z-position using a scaling factor of 0.67 ^76^; (2) setting the center-frequency ratio (cfr) to be above 0 (values below 0 are artifacts due to the freely floating imager strands in DNA-PAINT based measurements); (3) limiting the effective frequency at offset (efo) to detect only the first peak, representing single molecule events above background; (4) limiting the number of localizations (nLoc) above 5 to remove the unspecific binding events. The preprocessed data were then exported and converted to CSV files.

Subsequent analysis involved the following steps and was performed using the R programming language (R Core Team (2024). R: A Language and Environment for Statistical Computing. R Foundation for Statistical Computing, Vienna, Austria. https://www.R-project.org) with code available at https://git.embl.de/heriche/minflux-npc.

#### Particle extraction

A denoising step was applied to point clouds by removing points with fewer than 10 neighbours within a 20 nm radius. Particles were then isolated from the point cloud data using DBSCAN ^77^. Recovered structures were discarded if they contained fewer than 50 localization events or if the largest Feret’s diameter of their convex hull was below 20 nm or above 200 nm.

#### Clustering of particles

To distinguish ring-like structures from non-specific aggregates, extracted particles were further analyzed using topological data analysis followed by hierarchical clustering with Ward’s criterion, as previously described ^45^. The resulting dendrogram was cut to generate 2 clusters: Cluster 1, enriched in ring-like structures, and Cluster 2, containing singly labeled or irregular particles. Visual inspection of representative particles from each cluster confirmed the structural features identified by the clustering approach.

#### Diameter quantification

Ring-like NPCs were first registered using the joint registration of the multiple point cloud algorithm ^78^. Following mitosis, the reformed nuclear membrane has many folds and is therefore not always orthogonal to the imaging axis. To obtain a 2D projection of each NPC onto the nuclear membrane plane, registered NPCs were projected onto the plane orthogonal to the least axis of the globally registered NPCs. Diameters were then obtained by using the RANSAC algorithm ^79^ to fit a circle to a 2D projection of each structure along its least axis. Particles for which no robust circle fit could be obtained were excluded from further analysis.

### Sample preparation for correlative light-electron microscopy

The detailed protocol for sample preparation was published previously ^80^. In brief, cells were grown on carbon-coated sapphire disks (0.05 mm thick, 3 mm diameter; Wohlwend GmbH, Sennwald, Switzerland) in order to relocate cells of interest on EM grids. For orientation, a finder grid (Labtech, O7D00933) was overlaid on the sapphire disk during carbon evaporation.

Cell progression through mitotic exit was monitored by live imaging. Cells at different cell-cycle stages were then rapidly frozen using a high-pressure freezing machine (HPM 010; BalTec). Just prior to freezing, cells were immersed in imaging medium (IM) containing 20% Ficoll (Sigma Aldrich, PM400) to protect them from freezing-induced damage. It took ∼1 min from the last time-lapse imaging until the high-pressure freezing, and the time lag was recorded to precisely determine the duration after AO.

Freeze substitution was performed in a Leica EM AFS-2 freeze substitution unit (Leica Microsystems) as described previously ^80^. Briefly, the samples were substituted in 0.1% uranyl acetate (UA), 2% osmium tetroxide (Electron Microscopy Sciences, 19134) and 5% H_2_O in acetone (Electron Microscopy Sciences, cat 10015) following this temperature ramp: −90 °C to −80 °C for 10 h, −80 °C to −30 °C for 10 h, −30 °C for 4 h, −30 °C to 20 °C for 10 h, and 20 °C for 5-6 h. Afterwards, samples were washed three times in pure acetone for at least 10 min each, and subsequently infiltrated with Epon 812 hard formulation (Serva). The resin infiltration was done progressively at RT with increasing concentrations of resin in acetone (25% for 2-3 h, 50% for 2-3 h, and 75% 2-3h, 2 times 100% 1-2h plus 100% overnight). Resin was polymerized at 60 °C for 72 h.

Resin blocks were sectioned every 300 nm using a Diamond knife (Diatome) and an ultramicrotome Leica ultracut UC7 (Leica Microsystems). Sections were collected on Cu/Pd slot grids (Agar Scientific, G2564PD) coated with a film of 1% formvar (Agar Scientific) in chloroform (Sigma Aldrich, 32211-1L-M). To enhance membrane contrast, sections were post-stained at RT with 2% uranyl acetate in 70% methanol for 7 min and with 2% Reynold’s lead citrate in H2O (Delta Microscopies) for 4 min.

### Electron tomography (ET)

Single-axis tilt series were acquired with a TECNAI TF30 transmission EM (TEM; 300 kV; Thermo Fischer) equipped with a Gatan OneView 4k x 4k camera using the Serial EM software^81^. The samples were pre-irradiated by electron beam with the dose of 5-10 e/nm^2^ for at least 20 min to minimize sample shrinkage during tilt series acquisition. Images were recorded over a −60° to 60° tilt range with an angular increment of 1° at a pixel size of 1.0 nm. Tomograms were reconstructed using the R-weighted backprojection method implemented in the IMOD software package (version 4.5.6) ^81^.

### Membrane profile analysis

The nuclear membranes were manually traced by outlining them within the tomographic volume in the IMOD software package, as described previously ^3^. The ONM-INM distance and the pore diameter were measured from these 2D profiles using a custom analysis script written in MATLAB. This script was also used for membrane profile alignment and the calculation and display of the average profile. To generate the averaged NPC profile, each particle’s profile was aligned such that its symmetry axis is horizontal, dividing the cytoplasmic and nucleoplasmic halves, and centered at the same reference point. We then computed the mean intensity for the cytoplasmic and nucleoplasmic sides separately, sampling along the horizontal axis. Near the tip, we switch to polar sampling around the narrowing point so that the profile remains continuous. For the calculation of the ONM-INM distance, the median of the distance between 45 and 90 nm away from the edge of nuclear pores was measured.

### Quantitative blot analysis by Simple Western

To prepare the samples for Fig. 1. Suppl. 1B, total protein lysates were collected from cells grown in 10-cm dishes until ∼80% confluency. Cells were washed with PBS and resuspended in 500 μL of lysis buffer (RIPA buffer [Sigma-Aldrich, R0278], 1 mM PMSF [Sigma-Aldrich, P7626], cOmplete EDTA-free Protease Inhibitor Cocktail [Roche, 04693132001, 1 tablet/10 ml], and PhosSTOP [Roche, 4906845001, 1 tablet/10 ml]) on ice using a cell scraper.

To prepare the protein samples for Fig. 2 Suppl. 4 B-D, cells synchronized in prometaphase were harvested using the mitotic shake-off procedure ^14^. The resulting mitotic cell suspension was transferred to T-50 flasks and treated with either the dTAG compounds (250 nM of dTAG-13 and 500 nM of dTAG^V^-1) or the same amount of DMSO for 90 min in the presence of 82.5 nM nocodazole. Subsequently, the cells were collected into 15 ml Falcon tubes and centrifuged for 3 min at 90 × g. The cell pellets were resuspended in 1 mL of ice-cold PBS supplemented with cOmplete EDTA-free Protease Inhibitor Cocktail (1 tablet/10 ml), transferred to a 1.5 mL Eppendorf tube and centrifuged for 3 min at 90 × g and 4 °C. The resulting pellet was resuspended in 300 µL of ice-cold lysis buffer.

In both cases, cells were then lysed by two cycles of freezing in liquid nitrogen and thawing at 37°C. After centrifugation for 10 min at ∼16,000 × g and 4°C, the supernatants containing soluble protein extracts were collected and stored at −80°C until use.

Total protein concentration in cell extracts was measured using a Pierce BCA Protein Assay Kit (Thermo Fisher Scientific, 23227) and adjusted to 0.4 µg/µL by dilution in sample buffer (Bio-Techne, 042-195), including 1x Master Mix (from EZ Standard Pack 1 (Bio-Techne, PS-ST01EZ-8). Protein separation, immunodetection, and quantification from cell extracts were performed using a Jess Automated Western Blot System (Bio-Techne) with 12-230 kDa and 66-440 kDa fluorescence separation capillary cartridges (Bio-Techne, SM-FL004-1, SM-FL005-1). Protein samples and detection reagents were loaded into the Simple Western (SW) microplate following the manufacturer’s instructions. When protein normalization was required, the protein normalization module (Bio-Techne, DM-PN02) was used according to the manufacturer’s instructions. Detection was performed by ECL using anti-rabbit and anti-mouse secondary HRP and fluorescent antibodies (Bio-Techne, 042-206/042-205, 043-819/043-821) and Luminol-S/Peroxide solution (Bio-Techne, 043-311/043-379/043-8). Capillary electrophoresis was performed, and analysis was conducted using Compass for SW software (Bio-Techne), following the manufacturer’s guidelines.

### Molecular dynamics simulation of condensate formation by FG Nups

Three sets of simulations were used to study FG Nups. First, simulations of monomeric FG Nups were performed to reparametrize the force field for optimal modeling of FG Nups (*Training MOFF to accurately model FG Nups*). Second, slab simulations of individual FG Nup IDRs or the Nup62-Nup58-Nup54 subcomplex were performed to test the phase separation properties of FG Nups (*Slab simulations of FG Nups*). Third, constant volume simulations were performed to examine the surface tension and volume of Nup62-Nup58-Nup54 subcomplex condensates (*Constant concentration simulations of Nup62 subcomplexes*).

All simulations used a modified version of the MOFF force field ^52^, as described in *Training MOFF to accurately model FG Nups*. Additionally, 162 mM implicit, monovalent salt was used in all simulations, folded potentials were added to stabilize secondary structures and ordered motifs, and angles larger than 130° were rounded down to 130° to ensure numerical stability. All simulations were performed in OpenMM ^82^, with the OpenABC ^83^ implementation of MOFF, and are available at https://github.com/alatham13/Postmitotic_Nup62_OpenABC. All analysis code was implemented in MDAnalysis ^84^. Computational resources were provided by NSF’s Advanced Cyberinfrastructure Coordination Ecosystem: Services & Support (ACCESS) program and UCSF’s wynton high performance cluster ^85^.

#### Training MOFF to accurately model FG Nups

*T*o improve the force field accuracy for modeling FG Nups, we aimed to predict the *R_ee_* and *R_g_* of modified FG Nups (Nup49 [S. cerevisiae]: 121-154, Nup153 [H. sapiens]: 1313-1390, Nup153 [H. sapiens]: 884-993, Nup98 [H. sapiens]: 2-150, NSP1 [S. cerevisiae]: 2-175), which had previously been measured by Förster resonance energy transfer (FRET) and small-angle X-ray scattering (SAXS), respectively (exact sequences available in Table S2 of Fuertes et al.) ^86^. Forward predictions of these measurements were made in MOFF modified by a scaling coefficient (ε) with values 0.9, 0.95, 1.0, 1.05, or 1.1, where ε represents a small perturbation to the non-bonded potential of the MOFF force field of the form

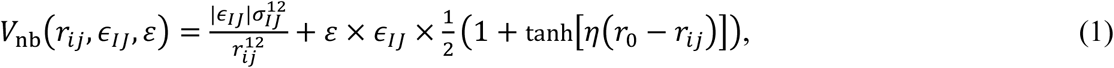

where *V*_nb_ is the strength of the non-bonded potential between amino residues, *r_ij_* is the distance between two amino residues, *ɛ_IJ_* is the strength of interactions between amino residues of type *I* and type *J*, *σ_IJ_* is the average size of amino residues of type *I* and type *J*, *η*=7 nm^-1^ is a smoothing parameter, and *r*_0_=0.8 nm is the distance cutoff of the contact potential.

Initial models for each protein in our training set were generated via ColabFold’s implementation of AlphaFold2 ^87,88^. Simulations used default parameters for MOFF besides ε (Eq. 1). Each protein was placed in a periodic simulation box with side lengths of 500 nm. We minimized the energy of the system, and then performed a 2 μs long constant-temperature simulation at 300 K using a Langevin integrator with a time coupling constant of 1 ps. Configurations were saved at 0.1 ns intervals and the first 1 μs was excluded for equilibration. Simulations were repeated five times with different random seeds for each protein.

After predicting the *R_ee_* and *R_g_* for each protein, we evaluated each set of simulations according to

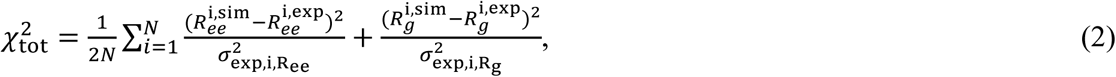

where *N* is the number of proteins in the training set, the sum is taken of each of the *i* proteins, 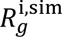 is the simulated *R_g_* for protein *i*, 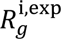 is the experimental *R_g_* for protein *i*, 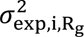 is the experimental standard deviation of the *R_g_* for protein *i*, 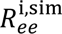 is the simulated *R_ee_* for protein *i*, 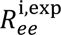 is the experimental *R_ee_* for protein *i*, and 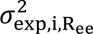 is the experimental estimate for the lower bound of precision of the *R_ee_* for protein *i*. Based on this oncriteria, ε=1.05 was selected for the remainder of our modeling (Figure 4 Supplementary 1B).

#### Slab simulations of FG Nups

To investigate the phase behavior of FG Nups, we performed direct-coexistence simulations ^89^ on each FG-repeat domain associated with Nup62 (Nup214 NT:1-699, Nup214 CT:973-2090, Nup62:1-331, Nup58 NT:1-245, Nup58 CT:419-599, Nup54:1-110) as well as the full length Nup62-Nup58-Nup54 subcomplex. Initial structures for the FG repeat domains were extracted from AlphaFold2 ^88^ structural predictions of the full length Nup, available from UniProt ^90^. The structure of the Nup62-Nup58-Nup54 subcomplex was predicted using ColabFold’s implementation of AlphaFold multimer ^87,91^, and tertiary contacts in the previously identified ^19^ ordered regions (i.e., the coiled-coil regions) were added to stabilize the trimer in MOFF. Simulations began with 100 copies of a specified Nup or subcomplex in a simulation box with 100 nm sides. This size has been shown to minimize finite-size effects ^92^. After energy minimization, constant-temperature, constant-pressure simulations were performed to condense to the protein in a single dense phase. This simulation lasted for 0.1 μs, was performed at 150 K and 1 bar, and used a Langevin integrator with a time coupling constant of 1 ps and a Monte Carlo barostat. Next, coexistence between the dense, protein phase, and dilute, solvent phase was created by expanding the Z-dimension of the simulation box to 500 nm. In this new simulation box, we raised the temperature of the simulation in a 0.1 μs constant-temperature simulation with a time coupling constant of 100 ps, where the target temperature was raised by 1.5 K every 1 ns, so that the simulation starts at 150 K and ends at 300 K. This starting point was run for production runs of 5 μs in the constant-temperature, constant-volume ensemble at 300 K. The first 2 μs were excluded for equilibration, and configurations were collected every 1 ns for analysis.

To analyze the slab simulations, we first computed a molecular contact matrix. Contacts between each subunit (either one copy of an FG Nup or one copy of the Nup62-Nup58-Nup54 subcomplex) were defined as any pair of subunits whose alpha-carbons were within 1 nm. Using a depth-first search algorithm over the network constructed from the contact matrix, we identified the largest cluster. The protein-density along the long axis (the z-dimension) was calculated relative to the center of mass of this largest cluster.

#### Constant concentration simulations of Nup62 subcomplexes

To quantify how the condensation properties of the Nup62-Nup58-Nup54 subcomplex may affect the assembly of the NPC, we performed constant number, temperature, and volume simulations with 32 copies of the Nup62-Nup58-Nup54 subcomplex to mirror the copy number in one NPC. We placed the 32 copies randomly within a square simulation box with side lengths of 81 nm, resulting in a Nup62-Nup58-Nup54 subcomplex concentration of 100 μM. Simulations lasted for 10 μs, used a Langevin thermostat with a time coupling constant of 100 ps, and had a target temperature of 300 K. The first 5 μs were excluded from analysis for equilibration, and configurations were collected every 1 ns for analysis.

Simulations were analyzed to determine the radius and surface tension of the droplet. Surface tensions were calculated according to the theory of Henderson and Lekner ^93^, which has previously been used to calculate the surface tension of FUS droplets from molecular dynamics simulations^94^. Briefly, the surface tension was estimated as

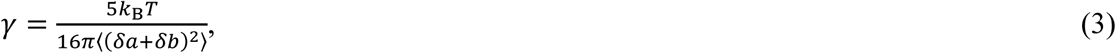

where *k*_B_ is Boltzmann’s constant, *T* is temperature, and *δa*=*a*-*R* as well as *δb*=*b*-*R* are the differences in lengths of any pair of principal axes of a general ellipsoid describing the instantaneous shape of the droplet with respect to the average radius of the droplet, *R*. Averages are taken over both all simulations times and all three principal axes. *R* was determined by fitting the radial mass density around the largest cluster, *c*(*r*), to a step function of the form

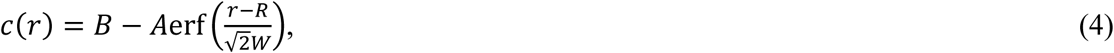

where *r* is the distance from the droplet center of mass, erf is the error function, and *A*, *B*, *R*, and *W* are determined by fitting.

Fifteen simulations beginning from unique starting configurations were performed and analyzed to determine R and γ. To account for inconsistencies in the quality of fit of *R* to Eq. 4, we removed simulations whose *R* estimates were more than three scaled median absolute deviations from the median. The reported R and γ are the average of the twelve remaining simulations. Volumes of the simulated condensates were calculated by assuming the condensate is a sphere of radius R.

This condensate volume was compared to the experimental volume of the nuclear pore before and after Nup62 depletion. For the experimental estimate, the pore was assumed to be a cylinder, whose height (36.82 nm for ΔNup62 and 28.16 nm for DMSO) and diameter (55.37 nm for ΔNup62 and 68.61 nm for DMSO) were given by ET data (Fig. 1C and E). We reported the difference in volume between DMSO and ΔNup62 conditions.

### Statistical analysis and sample size

All statistical analyses were performed using GraphPad Prism (version 10.5.0). For datasets following a normal distribution, comparisons between two groups were performed using two-tailed Welch’s t-tests, and comparisons involving more than two groups were analyzed using Welch’s ANOVA with Brown–Forsythe correction. For lognormally distributed data, values were log-transformed prior to analysis with either Welch’s t-tests (two groups) or Welch’s ANOVA (more than two groups). Non-parametric datasets were analyzed using two-tailed Mann–Whitney U tests (two groups) or Kruskal–Wallis tests followed by Dunn’s multiple comparisons test (more than two groups). No statistical methods were used to pre-determine sample sizes.

For EM tomography analysis, we analyzed 4-8 μm^2^ NE surface area per cell at different time points after acute perturbations. Here in the following are the details: for the acute Nup62 depletion assay, we analyzed cells divided at 20 and 24 min after AO and in interphase upon DMSO treatment, and at 20, 24, 28 and 34 min after AO upon dTAG treatment, yielding a total of 156 tomograms; for the HXD assay, we analyzed cells divided at 5, 6, 8 and 13 min after AO, as well as in interphase under HXD treatment and in interphase under control conditions, yielding 103 tomograms; for the acute import inhibition assay, we analyzed cells divided at 28 min after AO and in interphase upon TMP treatment, yielding 20 tomograms. For MINFLUX analysis, we analyzed cells at multiple time points upon different molecular perturbations. Here in the following are the details: for the acute Nup62 depletion assay, we analyzed cells divided at 21, 29, 32, 37 and 60 min after AO upon DMSO treatment, and at 20, 22, 24, 30, 31, 32, 42, 50 and 57 min after AO upon dTAG treatment; for the acute import inhibition assay, we analyzed cells under the control condition at 20, 21, 22, 30, 32, 37, 40 and 60 min after AO, and cells under the RanT24N-activitated condition at 20, 25, 30, 32, 37, 40 and 51 min after AO. For timelapse imaging, correlative immunofluorescence, and quantitative western blot analysis, data were obtained from at least two independent experiments. Sample sizes (n), measures of variation, and statistical significance are detailed in the figure legends.

## Supplementary Tables

**Supplementary Table 1:**
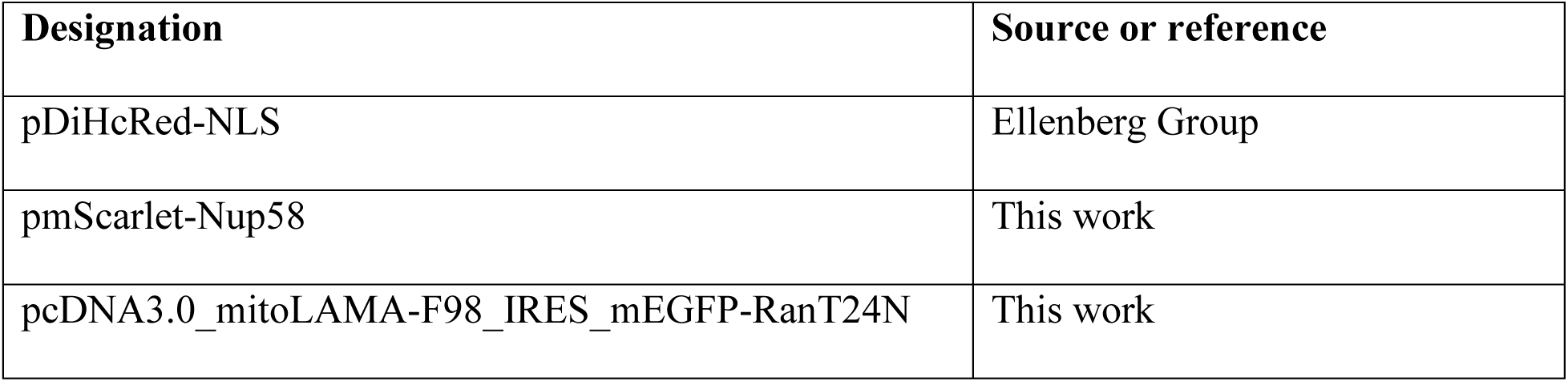
Plasmids used in this study.

**Supplementary Table 2:**
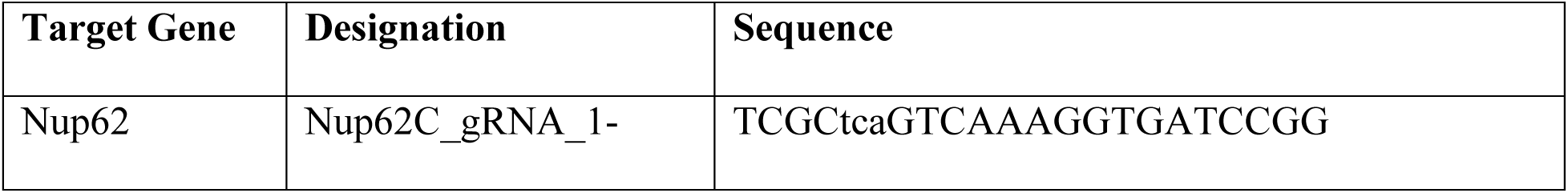
gRNA used in this study.

**Supplementary Table 3:** Antibodies used in this study.

| Antibodies | Dilution for<br>immune-<br>fluorescence | Dilution for<br>Simple Western | Source or reference |
| --- | --- | --- | --- |
| Anti-Elys | 1 to 50 |  | Protein Atlas Antibodies, HPA031658 |
| Anti-GFP |  | 1 to 50 | Abcam, ab290 |
| Anti-Nup107 |  | 1 to 50 | Proteintech, 19217-1-AP |
| Anti-Nup133 | 1 to 50 |  | Abcam, ab155990 |
| Anti-Nup153 | 1 to 500 |  | Abcam, ab96462 |
| Anti-Nup153 |  | 1 to 50 | Abcam, ab245668 |
| Anti-Nup155 | 1 to 100 |  | Protein Atlas Antibodies, HPA037775 |
| Anti-Nup188 |  | 1 to 50 | Thermo Fisher, A302-322A |
| Anti-Nup214 | 1 to 100 |  | Abcam, ab70497 |
| Anti-Nup214 |  | 1 to 50 | BETHYL, A300-716A |
| Anti-Nup358 | 1 to 100 |  | Protein Atlas Antibodies, HPA018437 |
| Anti-Nup54 |  | 1 to 25 | Protein Atlas Antibodies, HPA035929 |
| Anti-Nup58 |  | 1 to 25 | Invitrogen, PA5-113478 |
| Anti-Nup62 |  | 1 to 50 | BD Biosciences, 610497 |
| Anti-Nup88 | 1 to 50 | 1 to 25 | BD Transduction Laboratories™, 611896 |
| Anti-Nup98 | 1 to 50 |  | Cell Signaling Technology, 2598S |
| Anti-Tpr | 1 to 100 |  | Protein Atlas Antibodies, HPA019661 |
| Mab414 | 1 to 500 | 1 to 100 | BioLegend, 902902 |
| Anti-Rad21 | 1 to 500 |  | Merck Millipore, 05-908 |
| anti-Mouse-Abberior<br>STAR 635P | 1 to 250 |  | Abberior, ST635P-1001-500UG |
| anti-Rabbit-Abberior<br>STAR 580 | 1 to 250 |  | Abberior, ST580-1002-500UG |

**Supplementary Table 4:** Cell lines used in this study.

| Designation | Source or reference |
| --- | --- |
| HK WT | S. Narumiya (Kyoto University, Kyoto, Japan),<br>RRID: CVCL_1922 |
| HK Nup62-mEGFP-FKBP12<sup>F36V</sup> #02 | this work |
| HK Nup153-mEGFP-FKBP12<sup>F36V</sup> #10-043 | Brunner et al., 2025 |

## Supplementary Figures

**Figure 1_Supplementary 1.**
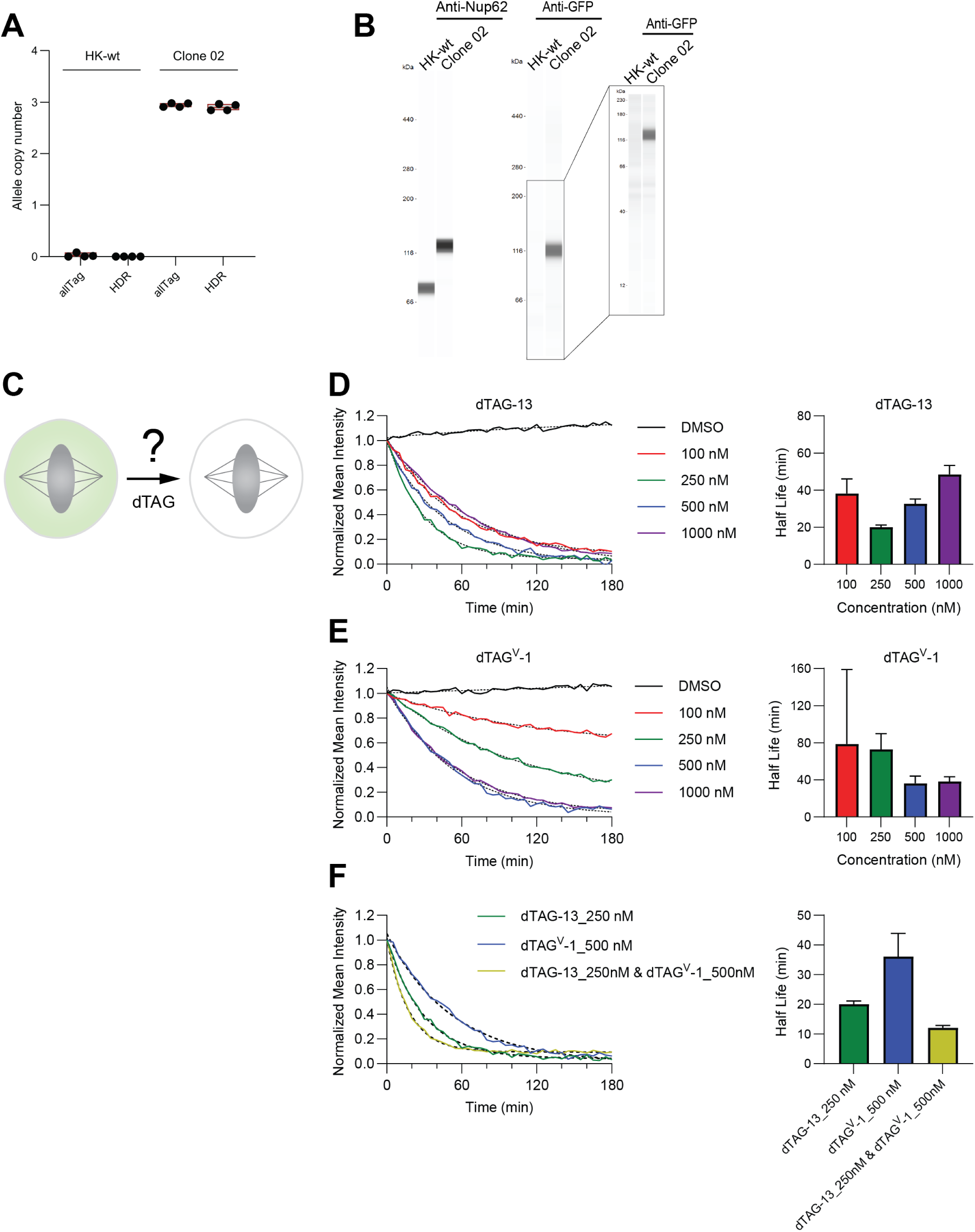
Validation of the homozygous Nup62-mEGFP-FKBP12^F36V^ knock-in cell line and optimization of acute Nup62 depletion during mitotic arrest. **(A, B**) Validation of the homozygous Nup62-mEGFP-FKBP12^F36V^ knock-in cell line (Clone 02). **(A)** Digital PCR analysis of homozygous integration of mEGFP-FKBP12^F36V^ at the Nup62 gene loci. Allele copy numbers of mEGFP-FKBP12^F36V^ integrations inserted into the HK genome (allTag) and at the endogenous Nup62 gene loci (HDR) were quantified from four replicates. Error bars indicate minimum and maximum values. **(B)** Simple Western analysis of total protein extracts from HK WT and the Nup62-mEGFP-FKBP12^F36V^ Clone 02 cells using antibodies against Nup62 and GFP. **(C)** Illustration of the assessment of Nup62-mEGFP-FKBP12^F36V^ degradation by dTAG compounds (dTAG-13 and/or dTAG^V^-1) during mitotic arrest. **(D, E)** Kinetics analysis of Nup62-mEGFP-FKBP12^F36V^ degradation by titrating dTAG-13 and dTAG^V^-1 concentrations. One-phase exponential decay models (dashed curves) were fitted to the data. Right panel: calculated half-life of degradation based on the fitted models. Sample sizes per condition: dTAG-13 (DMSO, n=13; 100 nM, n=7; 250 nM, n=9; 500 nM, n=12; 1000 nM, n=7) and dTAG^V^-1 (DMSO, n=9; 100 nM, n=11; 250 nM, n=8; 500 nM, n=6; 1000 nM, n=8). **(F)** Combination of 250 nM dTAG-13 and 500 nM dTAG^V^-1 (n=8) achieved the fastest degradation of Nup62-mEGFP-FKBP12^F36V^ among all tested conditions, as indicated by kinetic data and half-life calculated from fitted models (dashed curves).

**Figure 1_Supplementary 2.**
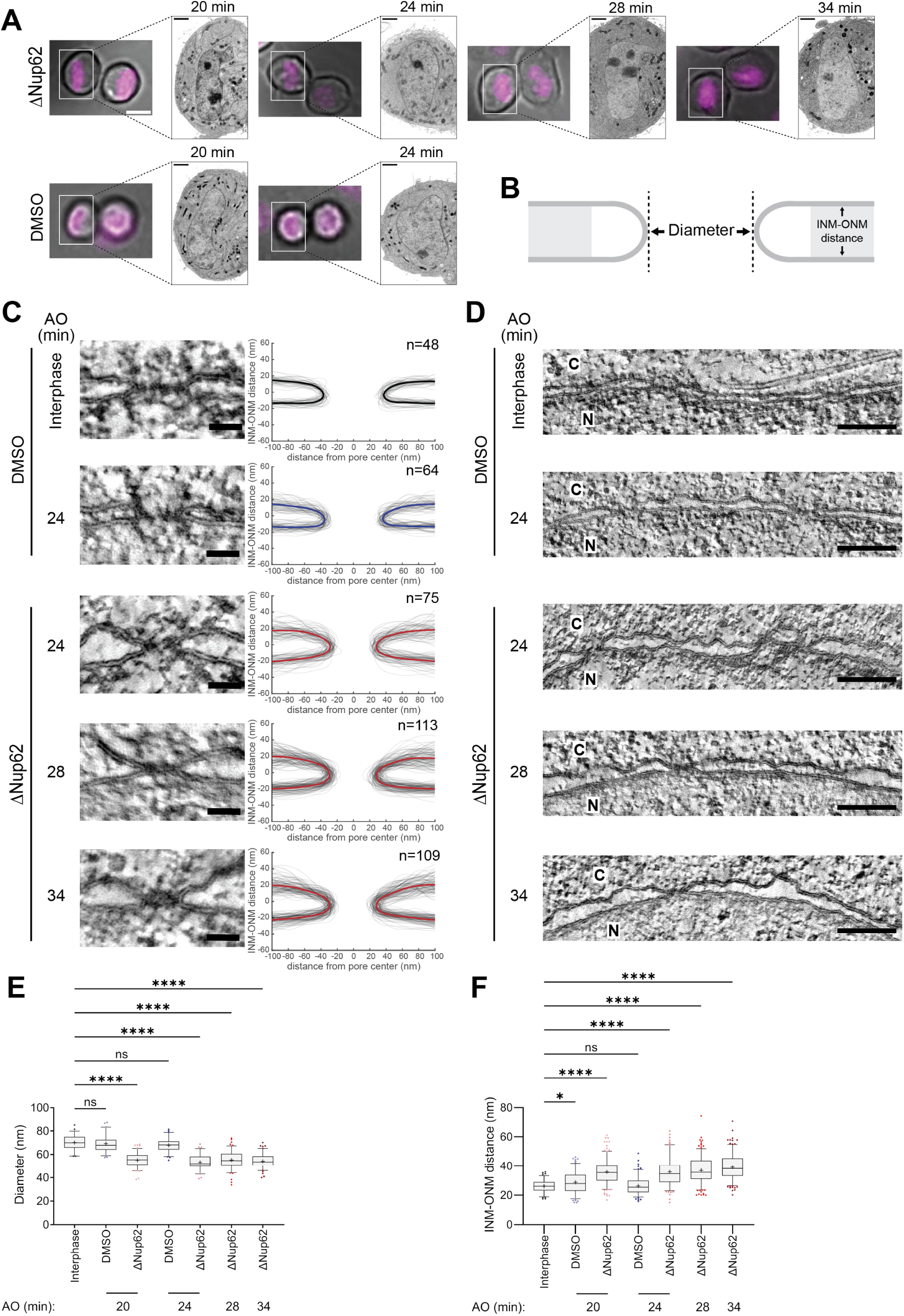
Nup62 depletion during mitosis reduces assembled pore diameter and increases NE spacing, persisting at later stages after AO. **(A)** Correlated live-cell and EM images of HK Nup62-mEGFP-FKBP^F36V^ cells analyzed by EM tomography. Cells treated with DMSO or dTAG (ΔNup62) were first imaged by light microscopy using live DNA dyes (shown in magenta) to track mitotic progression. The same cells were then subjected to high-pressure freezing, resin-embedding, serial sectioning, and observed in EM. Division time points after AO are indicated. Scale bars: light microscopy, 10 μm; EM, 2 μm. (**B)** Schematic illustrating how NPC diameter and INM–ONM distance were measured, indicated by bidirectional arrows. INM–ONM distance was calculated as the median of both sides of each measured pore within the indicated light gray areas (45-90 nm away from the pore tips). NPC diameter was measured as the distance between opposing membrane tips. **(C)** Left, tomographic slices showing cross-section views of nuclear pores from HK Nup62-mEGFP-FKBP^F36V^ cells treated with DMSO or dTAG (ΔNup62), at 24, 28, and 34 min after AO. NPCs from interphase cells treated with DMSO (> 180 min after AO) are shown for comparison. Right, membrane profiles of all measured pores at each time point are displayed, with mean profiles highlighted in bold. Scale bars, 50 nm. **(D)** Representative tomographic slices of the NE from the same samples as in (C). N, nucleus; C, cytoplasm. Scale bars, 200 nm. **(E, F)** Quantification of NPC diameter (E) and INM–ONM distance (F) from (C and D). Box-and-whisker plots show median, mean (“+”), 5–95 percentiles, and outliers (scatter). Sample sizes as indicated in (C) and Fig. 1B. Statistical significance applies to all panels in this figure: *, P ≤ 0.05; **, P ≤ 0.01; ***, P ≤ 0.001; ****, P ≤ 0.0001; ns, not significant.

**Figure 1_Supplementary 3.**
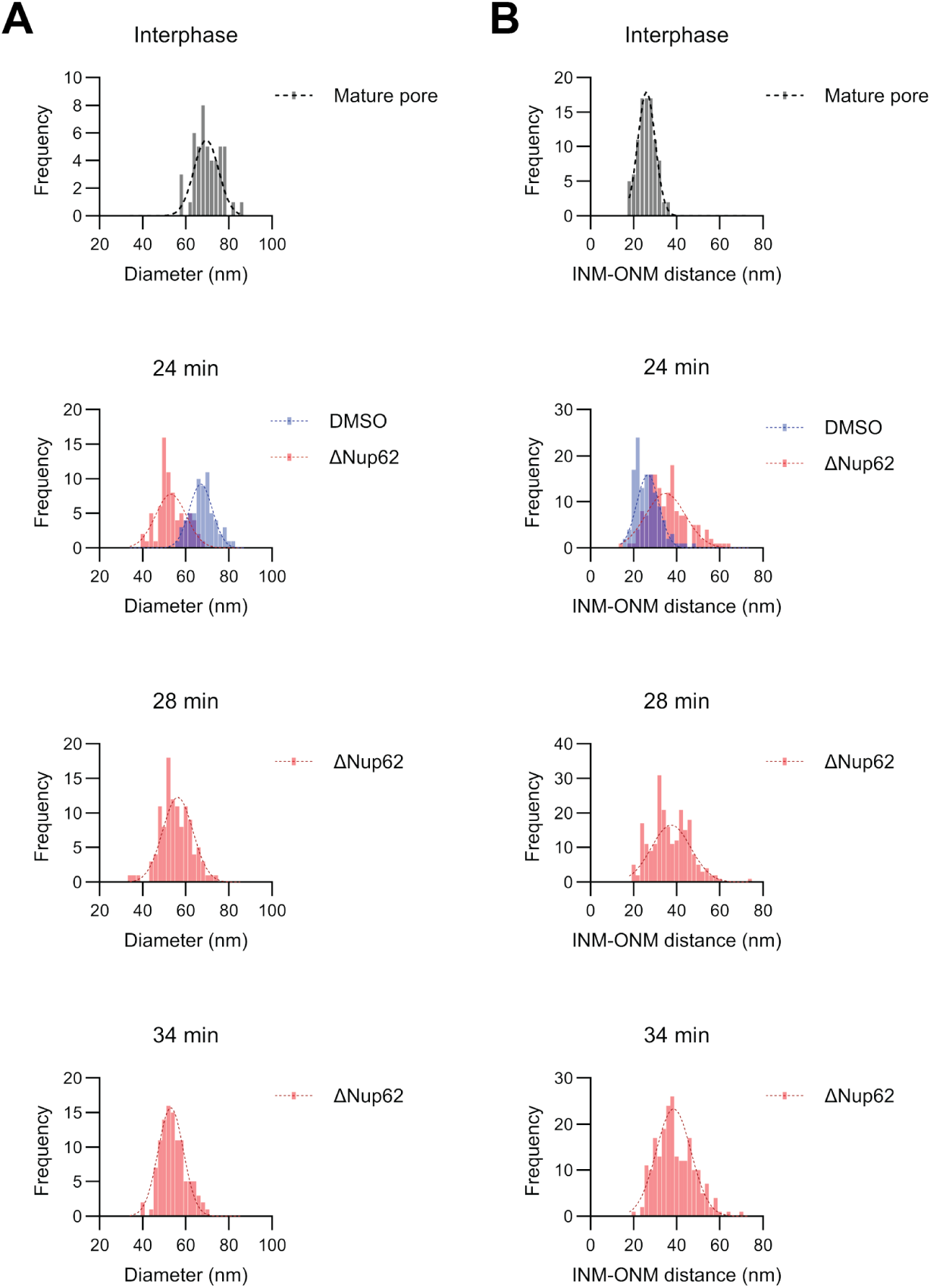
Distribution of NPC diameter and INM-ONM distance measured in Fig. 1 Suppl. 2E and F. **(A, B)** Histograms display the distribution of NPC diameter (A) and INM–ONM distance (B) at the indicated time points after AO, corresponding to Fig. 1 Suppl. 2E and F. Pores from interphase cells (> 180 min after AO) were analyzed for comparison (top panels). Gaussian distributions (dashed curves) were fitted to the histograms.

**Figure 1_Supplementary 4.**
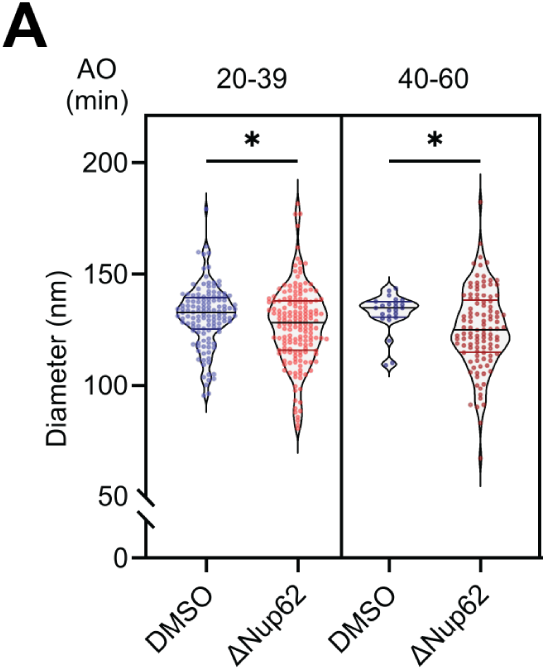
Nup62 depletion during mitosis reduces nuclear outer ring diameter of assembled NPCs. **(A)** Subgrouping of NPC diameter measurements according to the indicated time ranges after AO, corresponding to Fig. 1G. Median and quartiles are shown in the violin plot. Statistical significance applies to all panels in this figure: *, P ≤ 0.05; **, P ≤ 0.01; ***, P ≤ 0.001; ****, P ≤ 0.0001; ns, not significant.

**Figure 2_Supplementary 1.**
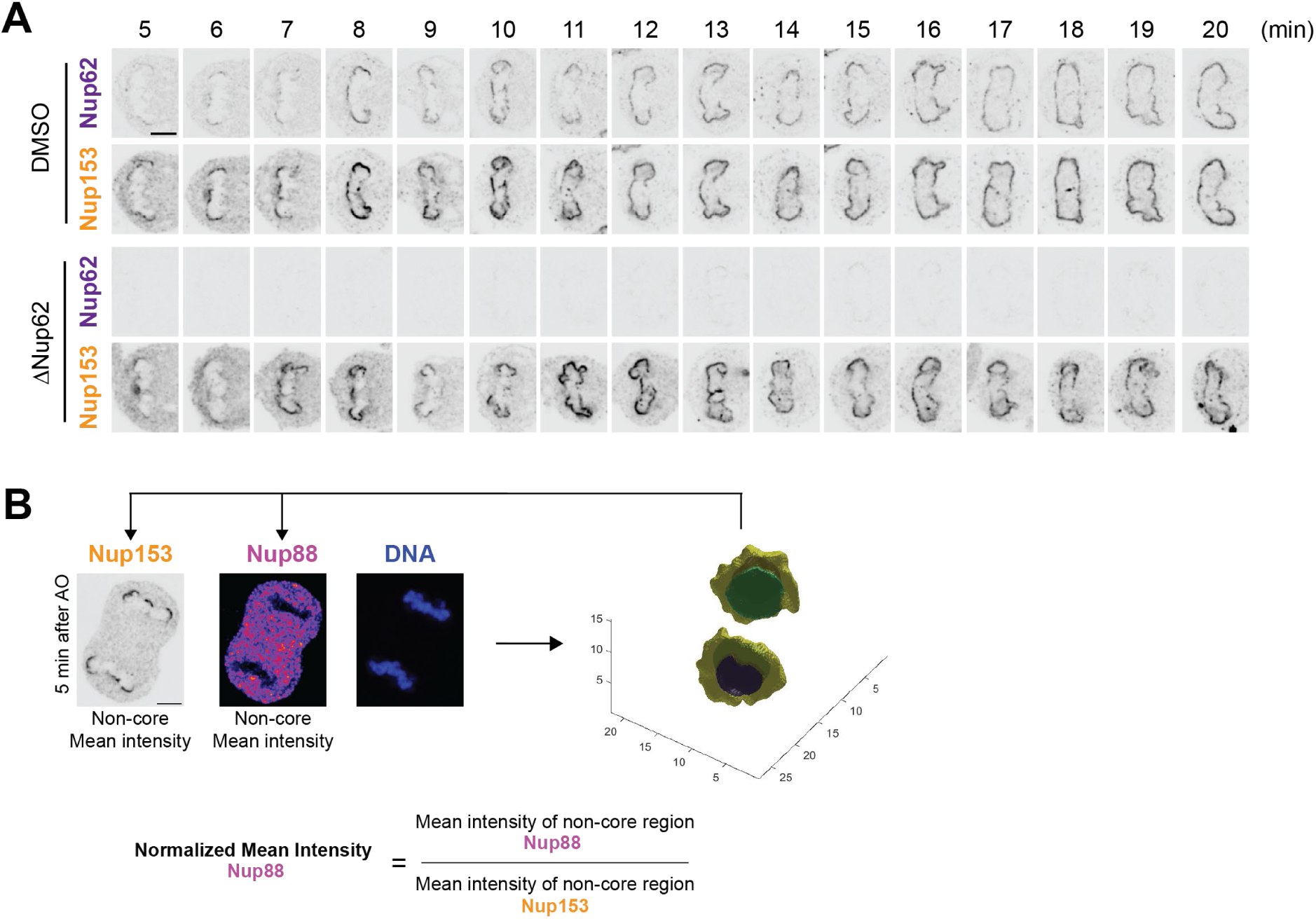
Loss of Nup62 does not affect the recruitment of Nup153 during mitotic exit and illustration of quantitative analysis used to normalize Nups recruitment by Nup153. **(A)** Representative images of HK Nup62-mEGFP-FKBP^F36V^ cells treated with DMSO or dTAG (ΔNup62), fixed at different time points (5-20 min) after AO, and stained with anti-Nup153 antibody. Nup62 signal is visualized via the mEGFP tagging. Scale bar, 5 μm. **(B)** Illustration of the normalized mean intensity calculation. A dividing cell at 5 min after AO, dually stained with anti-Nup153 and anti-Nup88 antibodies, is shown as an example. Scale bar, 5 μm. 3D segmented chromosomes based on the DNA channel and inferred non-core regions (yellow) are shown. Normalized mean intensity was calculated by dividing the measured mean intensity of Nup88 in non-core regions by the corresponding mean intensity of Nup153 measured in the same regions.

**Figure 2_Supplementary 2.**
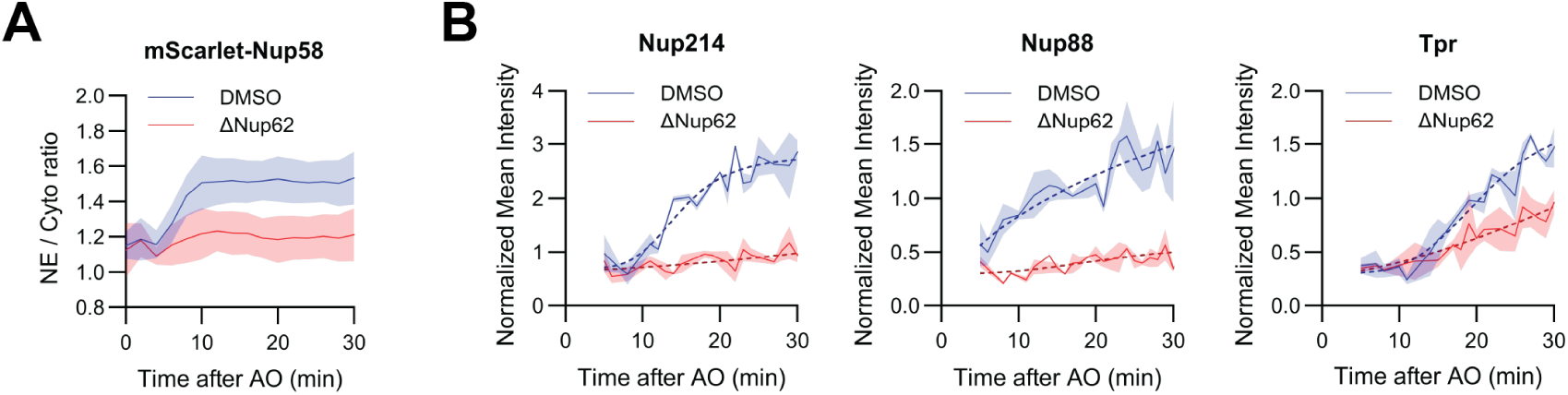
Loss of Nup62 impairs the recruitment of its interacting partners and the nuclear basket component. **(A)** Kinetic plot showing NE-to-cytoplasm mean intensity ratio of mScarlet-Nup58 in dividing HK Nup62-mEGFP-FKBP^F36V^ cells treated with DMSO or dTAG (ΔNup62) during mitotic exit (0-30 min after AO), corresponding to Fig. 2B. **(B)** Kinetic plots showing the normalized mean intensity of indicated Nups, measured from non-core regions of dividing HK Nup62-mEGFP-FKBP^F36V^ cells treated with DMSO or dTAG (ΔNup62), fixed at different time points (5-30 min) after AO, corresponding to Fig. 2C-E. Sigmoidal models (dashed curves) were fitted to the kinetic data.

**Figure 2_Supplementary 3.**
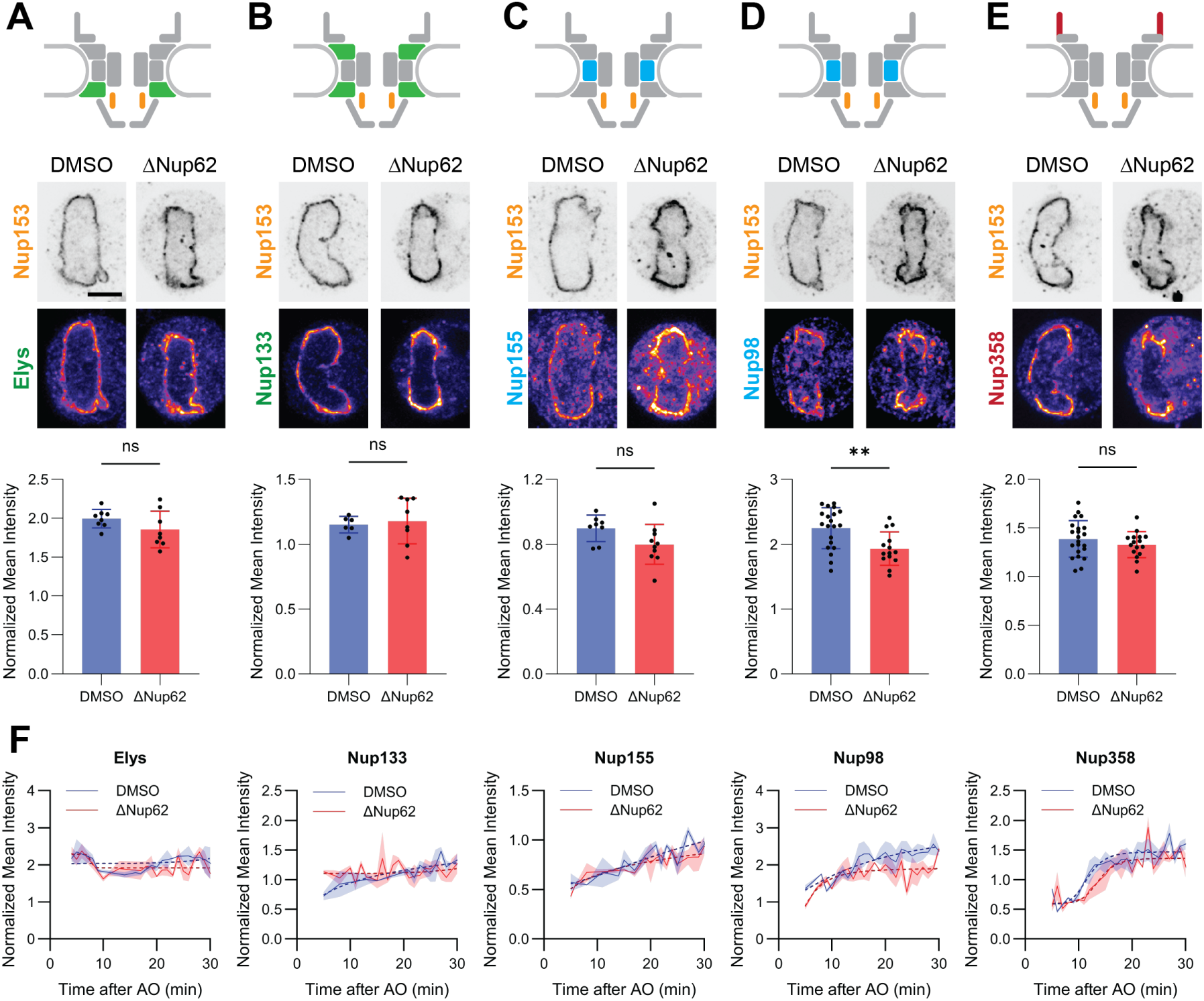
Loss of Nup62 does not affect the recruitment of scaffold Nups and the cytoplasmic filament. (A-E) Representative immunofluorescence images (middle panel) of HK Nup62-mEGFP-FKBP^F36V^ cells treated with DMSO or dTAG (ΔNup62) at 20 min after AO, dually stained with anti-Nup153 and anti-Elys (A), anti-Nup133 (B), anti-Nup155 (C), anti-Nup98 (D) or anti-Nup358 (E) antibodies. Upper panel: NPC schematic indicating the position of each labeled Nup within the complex. Lower panel: normalized mean intensity from non-core NE regions of dividing cells at 20 ± 1 min after AO. Scale bars, 5 μm. **(F)** Kinetic plots show the normalized mean intensity of indicated Nups shown in (C–G) calculated from non-core NE regions of cells fixed at different time points (5-30 min) after AO. Sigmoidal models (dashed curves) were fitted to the kinetic data. Statistical significance applies to all panels in this figure: *, P ≤ 0.05; **, P ≤ 0.01; ***, P ≤ 0.001; ****, P ≤ 0.0001; ns, not significant.

**Figure 2_Supplementary 4.**
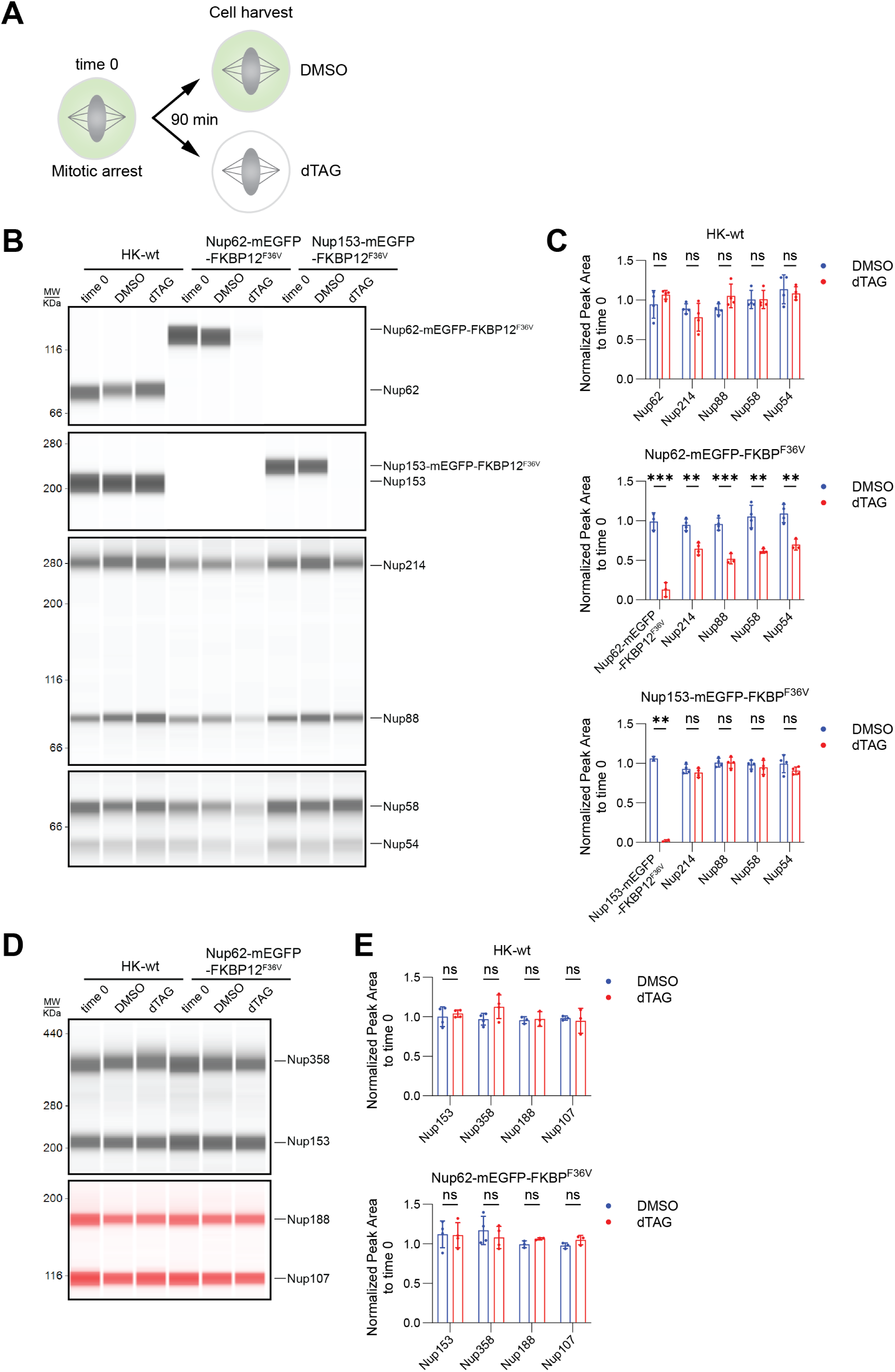
Nup62 depletion leads to a decrease in the protein levels of its interacting Nups. **(A)** Experimental setup for harvesting HK WT, HK Nup62-mEGFP-FKBP^F36V^ and HK Nup153-mEGFP-FKBP^F36V^ cells, arrested in prometaphase and treated with DMSO or dTAG (250 nM dTAG-13 and 500 nM dTAG^V^-1) for 90 min. **(B, D)** Simple Western analysis of protein extracts from the indicated cell lines treated as described in (A). For each condition, 3 μl of total protein lysate (0.4 µg/µl) was loaded into the microplates. The Jess ProteinSimple capillary system was used to detect total loaded proteins via the protein normalization module and specific Nups using targeted antibodies. In (B), antibodies were used in the following order (top to bottom panels): anti-Nup62, anti-Nup153, anti-Nup214 & anti-Nup88, and anti-Nup58 & anti-Nup54. In (D), antibodies used were anti-Mab414 (detecting Nup358 and Nup153) and anti-Nup188 & Nup107 (detected with secondary NIR antibodies). **(C, E)** Quantification of normalized protein levels of Nups detected in (B and D). Protein levels were first normalized to total protein content detected by the Jess ProteinSimple protein normalization module, and then further normalized to time 0 to assess relative changes. Statistical significance applies to all panels in this figure: *, P ≤ 0.05; **, P ≤ 0.01; ***, P ≤ 0.001; ****, P ≤ 0.0001; ns, not significant.

**Figure 3_Supplementary 1.**
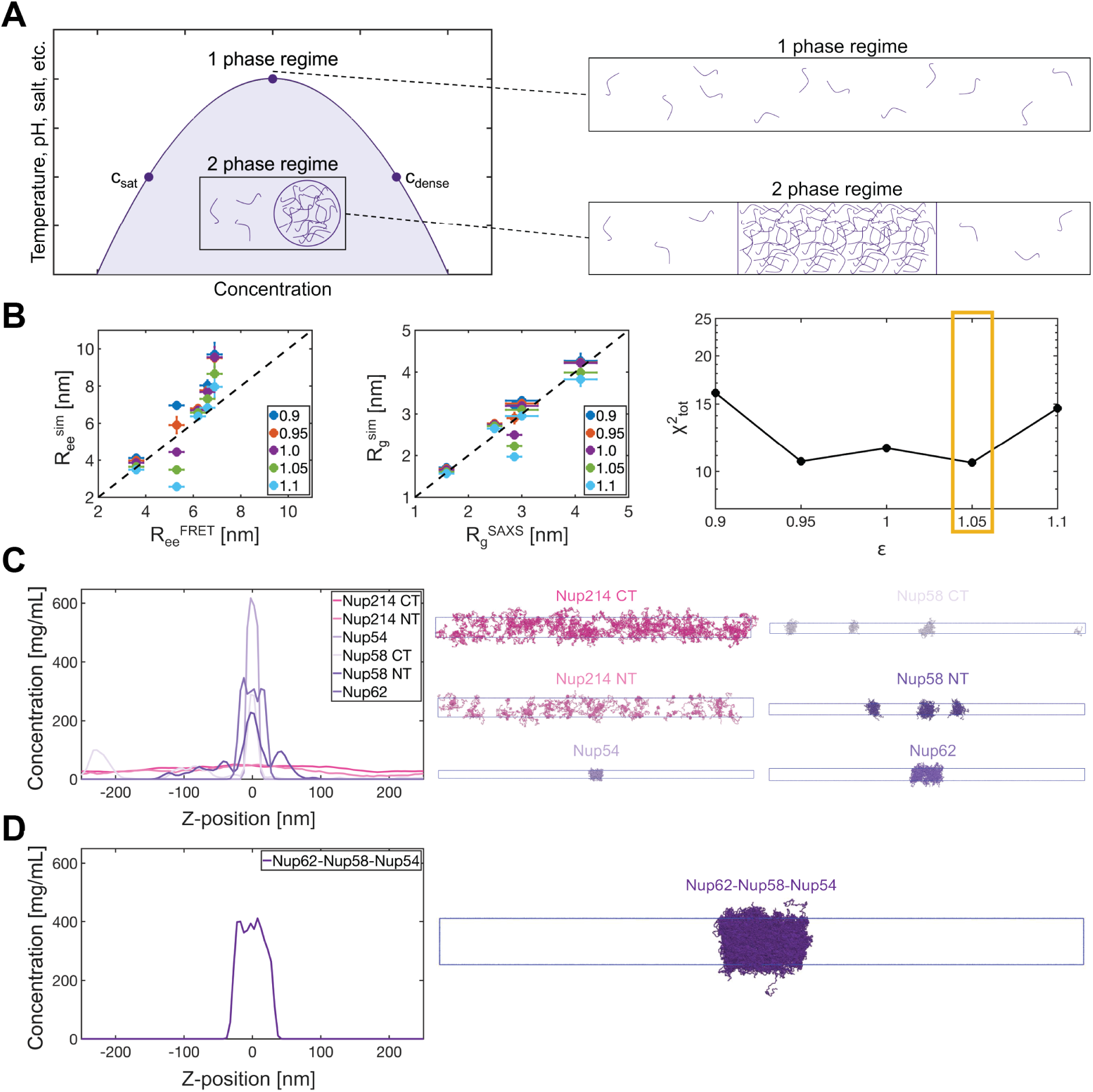
Simulation of condensate formation by Nup62-58-54 and Nup214-88-62 subcomplexes. **(A)** Illustration of a protein phase diagram. **(B)** Training MOFF to accurately model FG-Nups. Comparison between versions of MOFF with different ε to match experimental R_ee_ (left) or R_g_ (middle). Error bars in the X-axis represent standard deviations (SAXS) or the estimated lower bound of precision (FRET), while error bars in the Y-axis represent the standard deviation over five simulations with different random seeds. χ^2^ as a function of ε is also shown (right), with the optimal ε value used in modeling boxed. **(C)** Slab density profiles as a function of box length for IDRs of Nup214, Nup54, Nup58, and Nup62 (left), with images of the final frame from each simulation (right). **(D)** Slab density profile as a function of box length for the full length Nup62-Nup58-Nup54 subcomplex (left), with an image of the final frame from the simulation (right).

**Figure 3_Supplementary 2.**
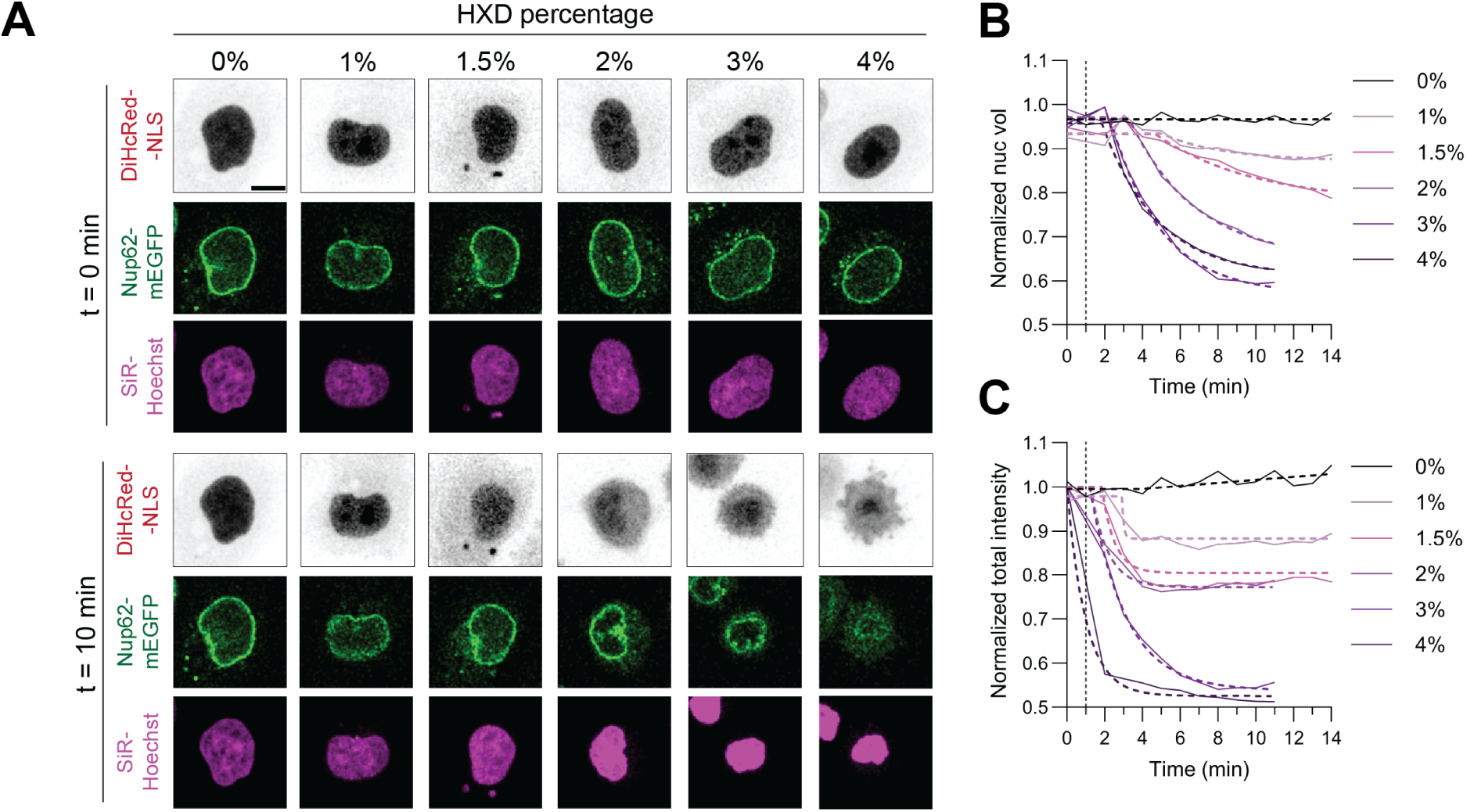
Acute 1,6-hexanediol (HXD) treatment affects nuclear volume and nuclear import in interphase. **(A)** HXD titration for acute disruption of FG-Nups condensate formation, assessed using the nuclear import reporter DiHcRed-NLS. HK Nup62-mEGFP-FKBP^F36V^ cells expressing DiHcRed-NLS and stained with SiR-Hoechst were imaged just before HXD addition (t = 0 min, upper panel) and after 10 min of treatment with the indicated HXD concentrations (lower panel). Scale bar, 10 μm. **(B, C)** Kinetic plots of nuclear volume (B) and total intensity of DiHcRed-NLS within the nucleus (C), normalized to t = 0, are shown over time. HXD was added at t = 1 min, as indicated by the dashed lines. Sample sizes per condition: 0%, n=14; 1%, n=14; 1.5%, n=10; 2%, n=12; 3%, n=11; 4%, n=12. Plateau followed by one-phase decay models (dashed curves) were fitted to the kinetic data.

**Figure 3_Supplementary 3.**
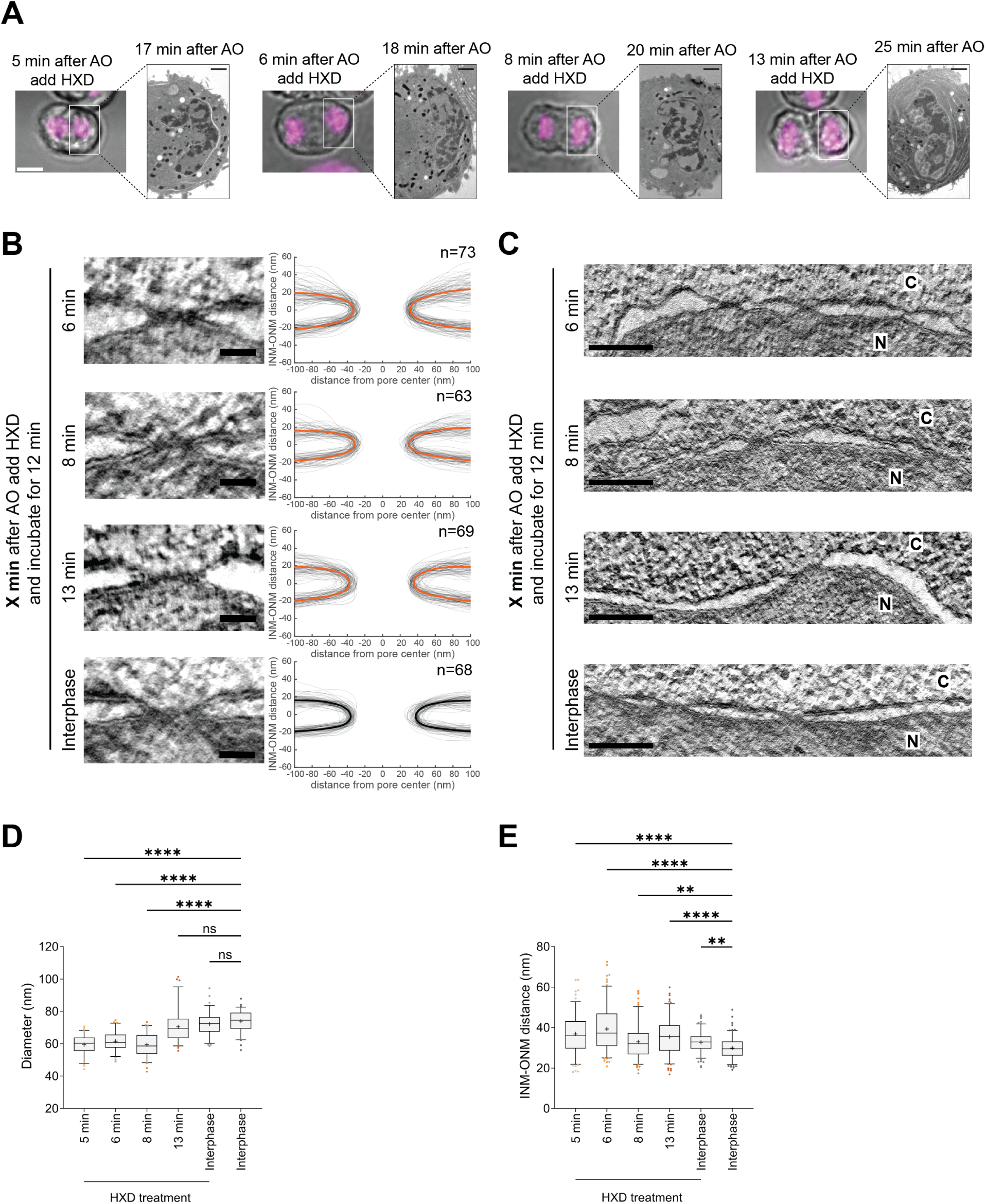
Acute disruption of condensate formation before pore dilation during NPC assembly reduces NPC diameter and increases NE spacing. **(A)** Correlated live-cell and EM images of HK WT cells analyzed by EM tomography. Cells were first imaged by light microscopy using live DNA dyes (shown in magenta) to track mitotic progression. Subsequently, 1.5% of HXD was added to cells at the indicated time points after AO and treated for 12 min. The same cells were then subjected to high-pressure freezing, resin-embedding, and serial sectioning, and observed in EM. Scale bars: light microscopy, 10 μm; EM, 2 μm. (**B)** Tomographic slices showing cross-section views of nuclear pores in HK WT cells treated with HXD at 6, 8, and 13 min after AO and in interphase. Membrane profiles of all measured pores are displayed, with mean profiles highlighted in bold. Scale bars, 50 nm. (**C)** Representative tomographic slices of the NE from the same samples as in (B). N, nucleus; C, cytoplasm. Scale bars, 200 nm. **(D, E)** Quantification of NPC diameter (D) and INM–ONM distance (E) from (C and D). Box-and-whisker plots show median, mean (“+”), 5–95 percentiles, and outliers (scatter). Sample sizes as indicated in (B) and Fig. 3E. Statistical significance applies to all panels in this figure: *, P ≤ 0.05; **, P ≤ 0.01; ***, P ≤ 0.001; ****, P ≤ 0.0001; ns, not significant.

**Figure 3_Supplementary 4.**
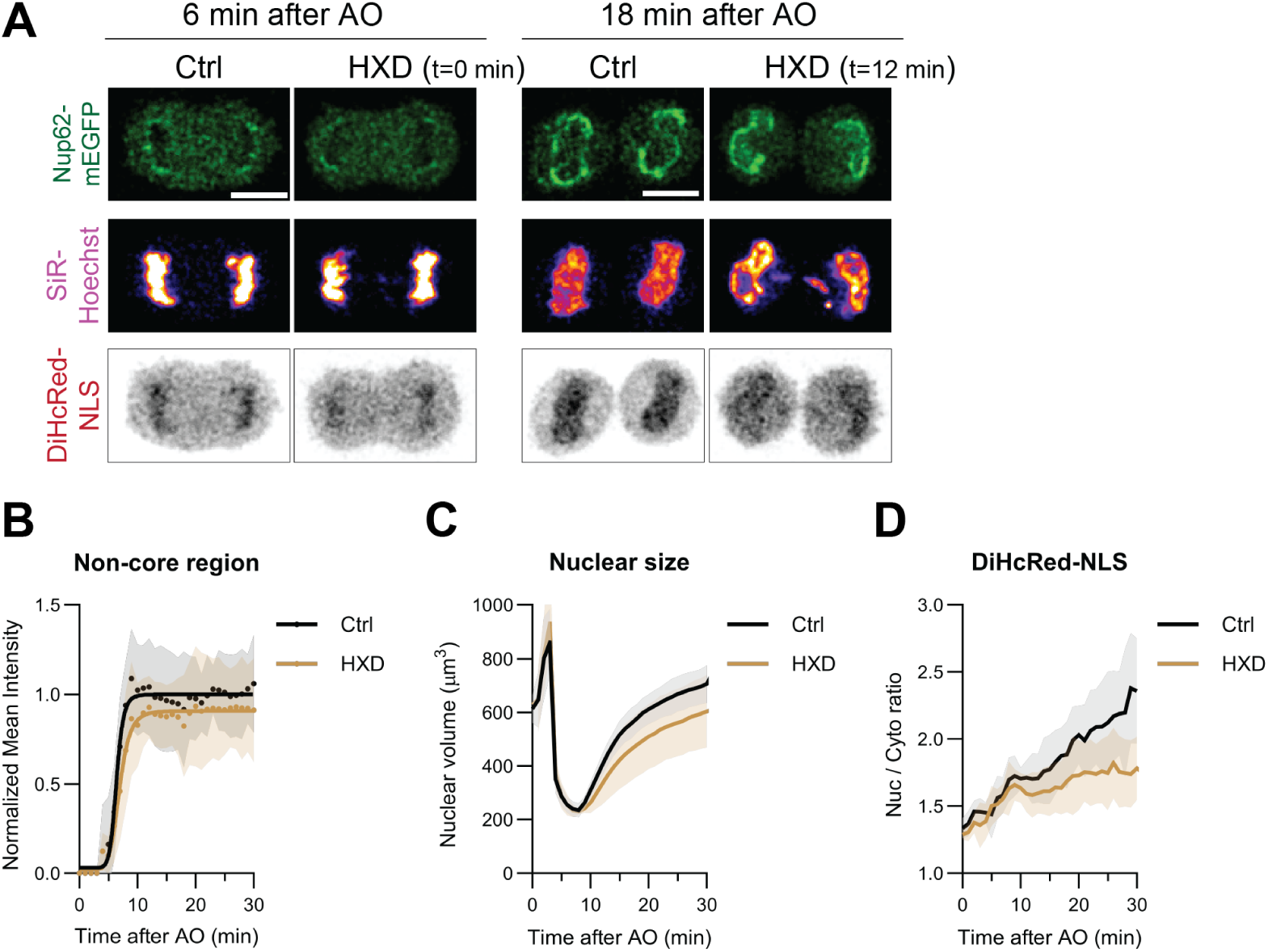
Acute 1,6-hexanediol (HXD) treatment affects nuclear volume and nuclear import during mitotic exit. **(A)** Representative live-cell images of HK Nup62-mEGFP-FKBP^F36V^ expressing DiHcRed-NLS at 6 min after AO, acutely treated with 1.5% of HXD for 12 min. Untreated cells served as controls. Scale bars, 10 μm. **(B)** Assembly kinetics of Nup62-mEGFP-FKBP^F36V^ in the non-core NE regions under control and HXD-treated conditions, from the same samples as in (A). Sigmoidal models (solid curves) were fitted to the kinetic data. Intensities were normalized to the plateau level of Nup62-mEGFP in the control condition. Data points represent the average normalized mean intensity, and error bands indicate the standard deviation. Sample sizes: Ctrl, n=16; HXD, n=12. **(C, D)** Kinetics of the average nuclear volume (C) and the nucleus-to-cytoplasm mean intensity ratio of DiHcRed-NLS (D) during mitotic exit, from the same samples as in (A). Error bands represent the standard deviation. Sample sizes: (C) Ctrl (n=16), HXD (n=12); (D) Ctrl (n=5), HXD (n=7).

**Figure 4_Supplementary 1.**
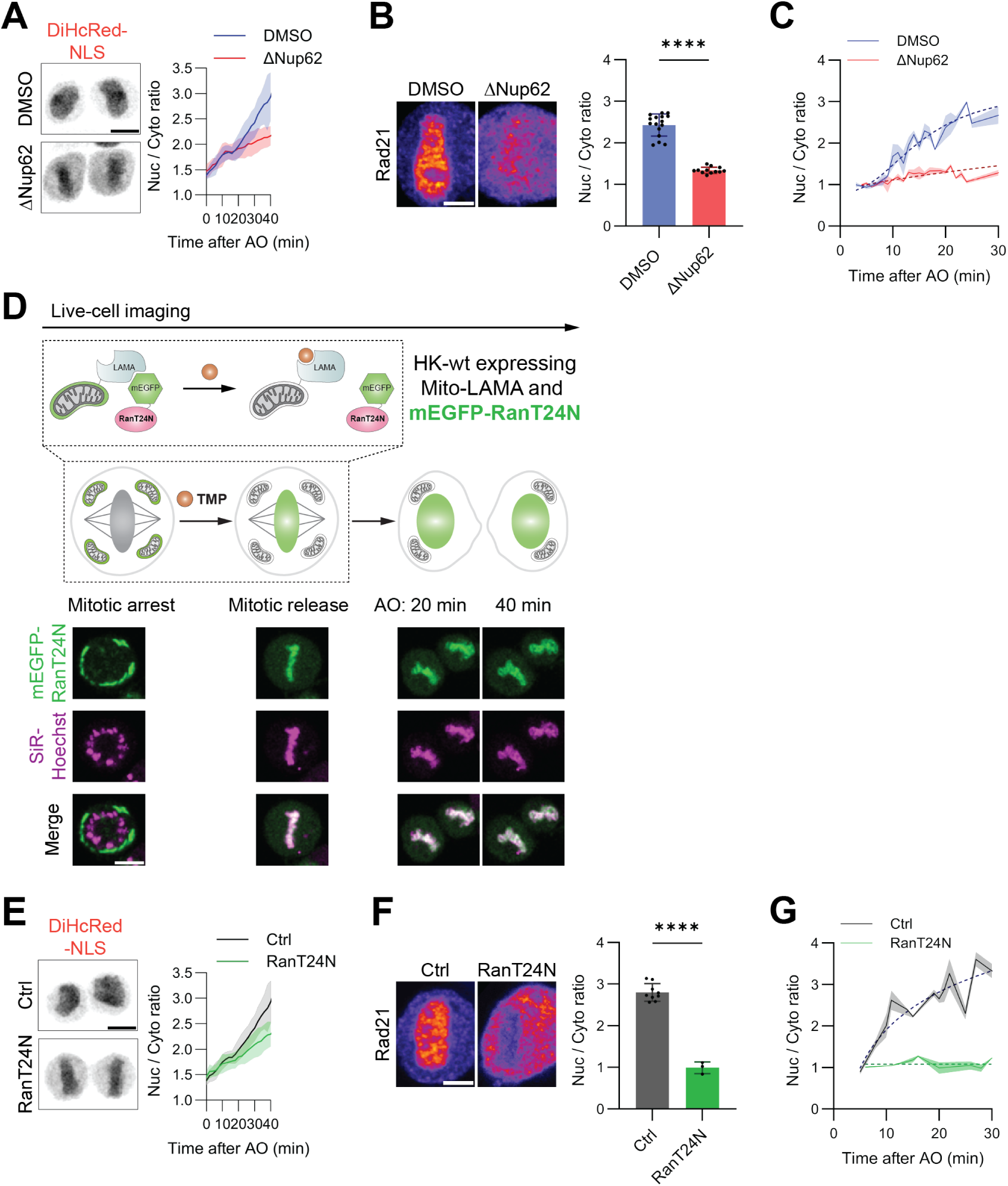
Nup62 depletion and RanT24N activation during mitotic exit impair nuclear import. **(A)** Representative live-cell images of HK Nup62-mEGFP-FKBP^F36V^ treated with DMSO or dTAG (ΔNup62), shown at 40 min after AO. Left, maximum projection of cells expressing DiHcRed-NLS. Scale bar, 10 μm. Right, nucleus-to-cytoplasm mean intensity ratio of DiHcRed-NLS measured in single cells during mitotic exit under the indicated conditions. Error bands represent the standard deviation. Sample sizes: DMSO, n=10; ΔNup62, n=12. **(B)** Representative correlative immunofluorescence images of HK Nup62-mEGFP-FKBP^F36V^ cells treated with DMSO or dTAG (ΔNup62), stained with anti-Rad21 at 20 min after AO. Scale bar, 5 μm. Right, nucleus-to-cytoplasm mean intensity ratio of Rad21 in dividing cells at 20 ± 1 min after AO. **(C)** Kinetic plot showing the nucleus-to-cytoplasm mean intensity ratio of Rad21 shown in (B), fixed at different time points (5-30 min) after AO. Sigmoidal models (dashed curves) were fitted to the kinetic data. (**D)** Schematic of the experimental workflow for acute import inhibition via transient release of mEGFP-RanT24N into the nucleus from mitochondria during mitotic exit using the mito-LAMA system. Release was triggered by adding 50 μM of TMP (illustrated in the upper panel). An example of a cell co-expressing mito-LAMA and mEGFP-RanT24N undergoing mitosis, after the release of mEGFP-RanT24N into the nucleus, corresponding to Fig. 4B. Scale bar, 10 μm. **(E)** Representative live-cell images of HK WT cells under Ctrl or RanT24N conditions, shown at 40 min after AO. Left, maximum projection of cells expressing DiHcRed-NLS; scale bar, 10 μm. Right, nucleus-to-cytoplasm mean intensity ratio of DiHcRed-NLS measured in single cells during mitotic exit under the indicated conditions. Error bands represent the standard deviation. Sample sizes: Ctrl, n=14; RanT24N, n=17. **(F)** Representative correlative immunofluorescence images of HK WT cells under Ctrl or RanT24N conditions, stained with anti-Rad21 at 20 min after AO. Scale bar, 5 μm. Right, nucleus-to-cytoplasm mean intensity ratio of Rad21 in dividing cells at 20 ± 1 min after AO. **(G)** Kinetic plot showing the nucleus-to-cytoplasm mean intensity ratio of Rad21 shown in (F), fixed at different time points (5-30 min) after AO. Sigmoidal models (dashed curves) were fitted to the kinetic data.

**Figure 4_Supplementary 2.**
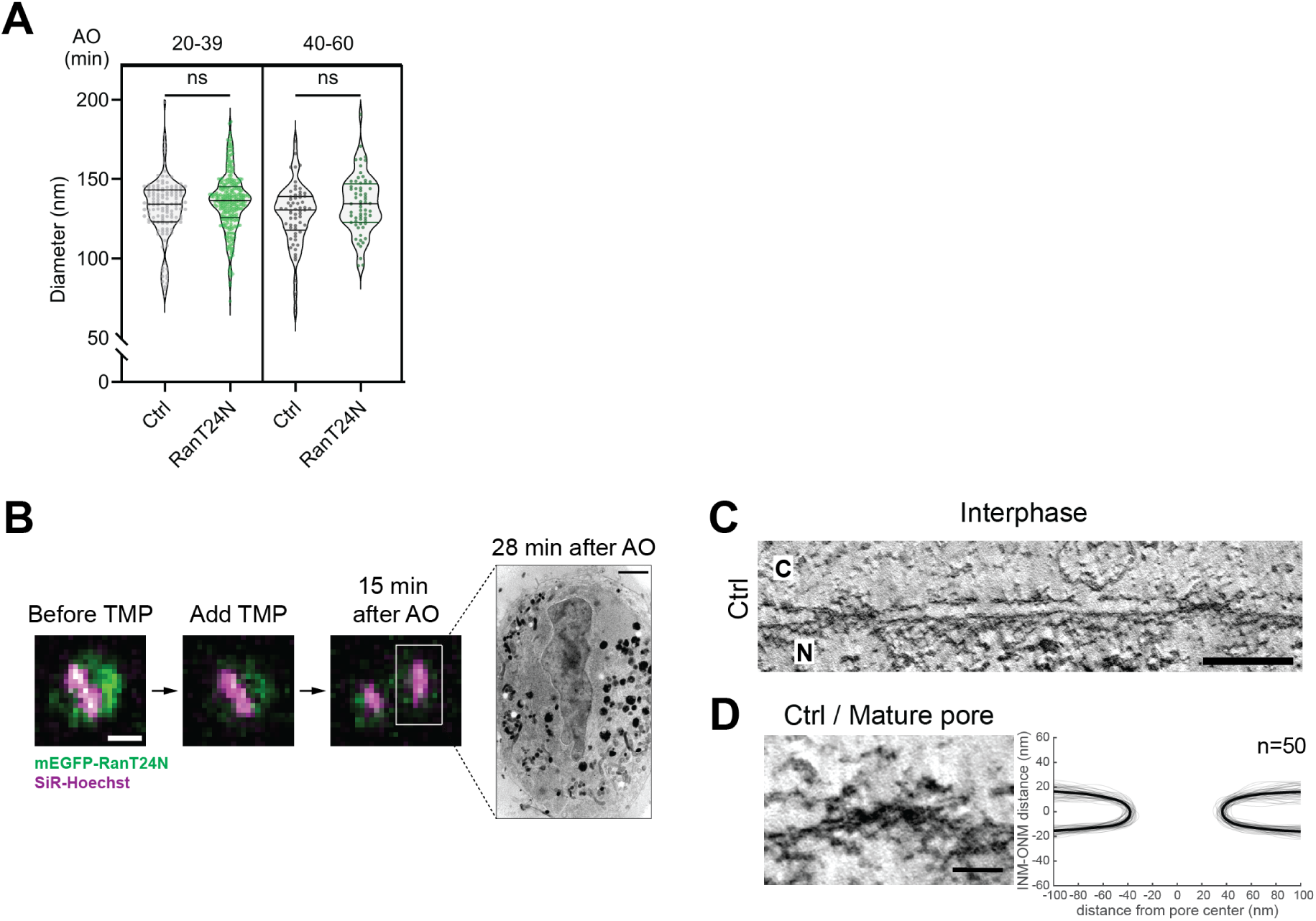
RanT24N activation during mitotic exit does not affect NPC diameter. **(A)** Subgrouping of pore diameter measurements according to the indicated division time ranges after AO, corresponding to Fig. 4E. Median and quartiles are shown in the violin plot. **(B)** Correlative live-cell and EM images of HK WT cells under RanT24N condition, analyzed by EM tomography. Cells were first imaged by light microscopy using live DNA dyes (shown in magenta) to track mitotic progression. Subsequently, 50 μM of TMP was then added to induce nuclear import of mEGFP-RanT24N during mitosis. The same cells were then subjected to high-pressure freezing, resin-embedding, serial sectioning, and observed in EM. Division time after AO is indicated. Scale bars: light microscopy, 10 μm; EM, 2 μm. **(C)** Representative tomographic slice of the NE in interphase HK WT cells under Ctrl condition. N, nucleus; C, cytoplasm. Scale bar, 200 nm. (**D)** Tomographic slices showing cross-section views of nuclear pores from the same samples as in (C). Membrane profiles of all measured pores are displayed, with mean profiles highlighted in bold. Scale bar, 50 nm. Statistical significance applies to all panels in this figure: *, P ≤ 0.05; **, P ≤ 0.01; ***, P ≤ 0.001; ****, P ≤ 0.0001; ns, not significant.

## References

1. Otsuka, S. & Ellenberg, J. Mechanisms of nuclear pore complex assembly – two different ways of building one molecular machine. FEBS Lett 592, 475–488 (2018).

2. Liu, S. & Pellman, D. The coordination of nuclear envelope assembly and chromosome segregation in metazoans. Nucleus 11, 35–52 (2020).

3. Otsuka, S. et al. Postmitotic nuclear pore assembly proceeds by radial dilation of small membrane openings. Nat Struct Mol Biol 25, 21–28 (2018).

4. Otsuka, S. et al. A quantitative map of nuclear pore assembly reveals two distinct mechanisms. Nature 613, 575–581 (2023).

5. Zhao, G. et al. A tubule-sheet continuum model for the mechanism of nuclear envelope assembly. Dev Cell 58, 847–865.e10 (2023).

6. Lu, L., Ladinsky, M. S. & Kirchhausen, T. Formation of the postmitotic nuclear envelope from extended ER cisternae precedes nuclear pore assembly. Journal of Cell Biology 194, 425–440 (2011).

7. Olmos, Y., Hodgson, L., Mantell, J., Verkade, P. & Carlton, J. G. ESCRT-III controls nuclear envelope reformation. Nature 522, 236–239 (2015).

8. Vietri, M. et al. Spastin and ESCRT-III coordinate mitotic spindle disassembly and nuclear envelope sealing. Nature 522, 231–235 (2015).

9. Kutay, U., Jühlen, R. & Antonin, W. Mitotic disassembly and reassembly of nuclear pore complexes. Trends in Cell Biology vol. xx 1–15 Preprint at 10.1016/j.tcb.2021.06.011 (2021).

10. Latham, A. P. et al. Integrative spatiotemporal modeling of biomolecular processes: Application to the assembly of the nuclear pore complex. Proceedings of the National Academy of Sciences 122, e2415674122 (2025).

11. Liu, S. et al. Nuclear envelope assembly defects link mitotic errors to chromothripsis. Nature 561, 551–555 (2018).

12. LaJoie, D. et al. A Role for Nup153 in Nuclear Assembly Reveals Differential Requirements for Targeting of Nuclear Envelope Constituents. Mol Biol Cell mbcE22050189 (2022) doi:10.1091/mbc.E22-05-0189.

13. Li, J., Jordana, L., Mehsen, H., Wang, X. & Archambault, V. Nuclear reassembly defects after mitosis trigger apoptotic and p53-dependent safeguard mechanisms in Drosophila. PLoS Biol 22, (2024).

14. Brunner, A. et al. Quantitative imaging of loop extruders rebuilding interphase genome architecture after mitosis. Journal of Cell Biology 224, e202405169 (2025).

15. Schooley, A. et al. Interphase chromosome conformation is specified by distinct folding programs inherited via mitotic chromosomes or through the cytoplasm. bioRxiv 2024.09.16.613305 (2024) doi:10.1101/2024.09.16.613305.

16. von Appen, A. et al. LEM2 phase separation promotes ESCRT-mediated nuclear envelope reformation. Nature 1–4 (2020) doi:10.1038/s41586-020-2232-x.

17. Dultz, E. et al. Systematic kinetic analysis of mitotic dis- and reassembly of the nuclear pore in living cells. Journal of Cell Biology 180, 857–865 (2008).

18. Anderson, D. J. & Hetzer, M. W. Reshaping of the endoplasmic reticulum limits the rate for nuclear envelope formation. Journal of Cell Biology 182, 911–924 (2008).

19. Mosalaganti, S. et al. AI-based structure prediction empowers integrative structural analysis of human nuclear pores. Science 376, eabm9506 (2022).

20. Bley, C. J. et al. Architecture of the cytoplasmic face of the nuclear pore. Science 1174, 1–66 (2022).

21. Petrovic, S. et al. Structure and Function of the Nuclear Pore Complex. Cold Spring Harb Perspect Biol (2022) doi:10.1101/cshperspect.a041264.

22. Petrovic, S. et al. Architecture of the linker-scaffold in the nuclear pore. Science 9798, 2021.10.26.465796 (2022).

23. Fontana, P. et al. Structure of cytoplasmic ring of nuclear pore complex by integrative cryo-EM and AlphaFold. Science 376, eabm9326 (2022).

24. Hamed, M., Caspar, B., Port, S. A. & Kehlenbach, R. H. A nuclear export sequence promotes CRM1-dependent targeting of the nucleoporin Nup214 to the nuclear pore complex. J Cell Sci 134, (2021).

25. Roth, P. et al. The Drosophila nucleoporin DNup88 localizes DNup214 and CRM1 on the nuclear envelope and attenuates NES-mediated nuclear export. Journal of Cell Biology 163, 701–706 (2003).

26. Bernad, R., Engelsma, D., Sanderson, H., Pickersgill, H. & Fornerod, M. Nup214-Nup88 Nucleoporin Subcomplex Is Required for CRM1-mediated 60 S Preribosomal Nuclear Export*. Journal of Biological Chemistry 281, 19378–19386 (2006).

27. Cowburn, D. & Rout, M. Improving the hole picture: towards a consensus on the mechanism of nuclear transport. Biochem Soc Trans 51, 871–886 (2023).

28. Kehlenbach, R. H., Neumann, P., Ficner, R. & Dickmanns, A. Interaction of nucleoporins with nuclear transport receptors: a structural perspective. Biol Chem 404, 791–805 (2023).

29. Aramburu, I. V. & Lemke, E. A. Floppy but not sloppy: Interaction mechanism of FG-nucleoporins and nuclear transport receptors. Semin Cell Dev Biol 68, 34–41 (2017).

30. Ng, S. C. & Görlich, D. A simple thermodynamic description of phase separation of Nup98 FG domains. Nat Commun 13, 1–17 (2022).

31. Dekker, M., Van der Giessen, E. & Onck, P. R. Phase separation of intrinsically disordered FG-Nups is driven by highly dynamic FG motifs. Proceedings of the National Academy of Sciences 120, e2221804120 (2023).

32. Nag, N., Sasidharan, S., Uversky, V. N., Saudagar, P. & Tripathi, T. Phase separation of FG-nucleoporins in nuclear pore complexes. Biochimica et Biophysica Acta (BBA) - Molecular Cell Research 1869, 119205 (2022).

33. Schmidt, H. B. & Görlich, D. Nup98 FG domains from diverse species spontaneously phase-separate into particles with nuclear pore-like permselectivity. Elife 4, e04251 (2015).

34. Ng, S. C. et al. Barrier properties of Nup98 FG phases ruled by FG motif identity and inter-FG spacer length. Nat Commun 14, 747 (2023).

35. Ibáñez de Opakua, A., Pantoja, C. F., Cima-Omori, M. S., Dienemann, C. & Zweckstetter, M. Impact of distinct FG nucleoporin repeats on Nup98 self-association. Nat Commun 15, (2024).

36. Celetti, G. et al. The liquid state of FG-nucleoporins mimics permeability barrier properties of nuclear pore complexes. Journal of Cell Biology 219, (2020).

37. Patel, S. S., Belmont, B. J., Sante, J. M. & Rexach, M. F. Natively Unfolded Nucleoporins Gate Protein Diffusion across the Nuclear Pore Complex. Cell 129, 83–96 (2007).

38. Raveh, B., et al. Integrative mapping reveals molecular features underlying the mechanism of nucleocytoplasmic transport. bioRxiv 2023.12.31.573409 (2025) doi:10.1101/2023.12.31.573409.

39. Timney, B. L. et al. Simple rules for passive diffusion through the nuclear pore complex. Journal of Cell Biology 215, 57–76 (2016).

40. Delavoie, F., Soldan, V., Rinaldi, D., Dauxois, J. Y. & Gleizes, P. E. The path of pre-ribosomes through the nuclear pore complex revealed by electron tomography. Nat Commun 10, 1–12 (2019).

41. Jevtić, P. et al. The nucleoporin ELYS regulates nuclear size by controlling NPC number and nuclear import capacity. EMBO Rep 20, 1–16 (2019).

42. Callegari, A., Kueblbeck, M., Morero, N. R., Serrano-Solano, B. & Ellenberg, J. Rapid generation of homozygous fluorescent knock-in human cells using CRISPR–Cas9 genome editing and validation by automated imaging and digital PCR screening. Nat Protoc 20, 26–66 (2025).

43. Nabet, B. et al. The dTAG system for immediate and target-specific protein degradation. Nat Chem Biol 14, 431–441 (2018).

44. Nabet, B. et al. Rapid and direct control of target protein levels with VHL-recruiting dTAG molecules. Nat Commun 11, (2020).

45. Sabinina, V. J. et al. Three-dimensional superresolution fluorescence microscopy maps the variable molecular architecture of the nuclear pore complex. Mol Biol Cell 32, 1523–1533 (2021).

46. Hoffmann, P. C. et al. Nuclear pore permeability and fluid flow are modulated by its dilation state. Mol Cell (2024) doi:10.1016/j.molcel.2024.11.038.

47. Zimmerli, C. E. et al. Nuclear pores dilate and constrict in cellulo. Science 9776, eabd9776 (2021).

48. Heusel, M. et al. A Global Screen for Assembly State Changes of the Mitotic Proteome by SEC-SWATH- MS. Cell Syst 10, 133–155.e6 (2020).

49. Linder, M. I. et al. Mitotic Disassembly of Nuclear Pore Complexes Involves CDK1- and PLK1-Mediated Phosphorylation of Key Interconnecting Nucleoporins. Dev Cell 43, 141–156.e7 (2017).

50. Nkoula, S. N. et al. Mechanisms of nuclear pore complex disassembly by the mitotic Polo-like kinase 1 (PLK-1) in *C. elegans* embryos. Sci Adv 9, eadf7826 (2023).

51. Wang, J. et al. Phase separation of the nuclear pore complex facilitates selective nuclear transport to regulate plant defense against pathogen and pest invasion. Mol Plant 16, 1016–1030 (2023).

52. Latham, A. P. & Zhang, B. Consistent Force Field Captures Homologue-Resolved HP1 Phase Separation. J Chem Theory Comput 17, 3134–3144 (2021).

53. Mangiarotti, A., Chen, N., Zhao, Z., Lipowsky, R. & Dimova, R. Wetting and complex remodeling of membranes by biomolecular condensates. Nat Commun 14, 1–15 (2023).

54. Mondal, S. & Baumgart, T. Membrane reshaping by protein condensates. Biochim Biophys Acta Biomembr 1865, 184121 (2023).

55. Barrientos, E. C. R., Otto, T. A., Mouton, S. N., Steen, A. & Veenhoff, L. M. A survey of the specificity and mechanism of 1,6 hexanediol-induced disruption of nuclear transport. Nucleus 14, 2240139 (2023).

56. Farrants, H. et al. Chemogenetic Control of Nanobodies. Nat Methods 17, 279–282 (2020).

57. Ben-Efraim, I., Frosst, P. D. & Gerace, L. Karyopherin binding interactions and nuclear import mechanism of nuclear pore complex protein Tpr. BMC Cell Biol 10, 74 (2009).

58. Snow, C. J., Dar, A., Dutta, A., Kehlenbach, R. H. & Paschal, B. M. Defective nuclear import of Tpr in Progeria reflects the Ran sensitivity of large cargo transport. Journal of Cell Biology 201, 541–557 (2013).

59. Huang, G. et al. Structure of the Cytoplasmic Ring of the Xenopus laevis Nuclear Pore Complex. Science., 2020.03.27.009407 (2022).

60. Gouveia, B. et al. Capillary forces generated by biomolecular condensates. Nature 609, 255–264 (2022).

61. Yuan, F. et al. Membrane bending by protein phase separation. Proceedings of the National Academy of Sciences 118, e2017435118 (2021).

62. Wang, Y. et al. Biomolecular condensates mediate bending and scission of endosome membranes. Nature (2024) doi:10.1038/s41586-024-07990-0.

63. Pombo-García, K., Adame-Arana, O., Martin-Lemaitre, C., Jülicher, F. & Honigmann, A. Membrane prewetting by condensates promotes tight-junction belt formation. Nature 632, 647–655 (2024).

64. Mangiarotti, A. & Dimova, R. Biomolecular Condensates in Contact with Membranes. Annu Rev Biophys 53, 319–341 (2024).

65. Kim, N., Yun, H., Lee, H. & Yoo, J. Y. Interplay between membranes and biomolecular condensates in the regulation of membrane-associated cellular processes. Exp Mol Med 34, (2024).

66. Shulga, N. & Goldfarb, D. S. Binding Dynamics of Structural Nucleoporins Govern Nuclear Pore Complex Permeability and May Mediate Channel Gating. Mol Cell Biol 23, 534–542 (2003).

67. Mauro, M. S. et al. Ndc1 drives nuclear pore complex assembly independent of membrane biogenesis to promote nuclear formation and growth. Elife 11, (2022).

68. Agrawal, A. & Lele, T. P. Mechanics of nuclear membranes. J Cell Sci 132, (2019).

69. Torbati, M., Lele, T. P. & Agrawal, A. Ultradonut topology of the nuclear envelope. Proceedings of the National Academy of Sciences 113, 11094–11099 (2016).

70. Taniguchi, R. et al. Nuclear pores safeguard the integrity of the nuclear envelope. Nat Cell Biol (2025) doi:10.1038/s41556-025-01648-3.

71. Elosegui-Artola, A. et al. Force Triggers YAP Nuclear Entry by Regulating Transport across Nuclear Pores. Cell 171, 1397–1410.e14 (2017).

72. Otsuka, S. et al. Nuclear pore assembly proceeds by an inside-out extrusion of the nuclear envelope. Elife 5, 1–23 (2016).

73. Bucevičius, J., Keller-Findeisen, J., Gilat, T., Hell, S. W. & Lukinavičius, G. Rhodamine–Hoechst positional isomers for highly efficient staining of heterochromatin. Chem Sci 10, 1962–1970 (2019).

74. Balzarotti, F. et al. Nanometer resolution imaging and tracking of fluorescent molecules with minimal photon fluxes. Science (1979) 355, 606–612 (2017).

75. Schmidt, R. et al. MINFLUX nanometer-scale 3D imaging and microsecond-range tracking on a common fluorescence microscope. Nat Commun 12, 1478 (2021).

76. Sau, A. et al. Overlapping nuclear import and export paths unveiled by two-colour MINFLUX. Nature (2025) doi:10.1038/s41586-025-08738-0.

77. Ester, M., Kriegel, H.-P., Sander, J. & Xu, X. A density-based algorithm for discovering clusters in large spatial databases with noise. in Proceedings of the Second International Conference on Knowledge Discovery and Data Mining 226–231 (AAAI Press, 1996).

78. Evangelidis, G. D., Kounades-Bastian, D., Horaud, R. & Psarakis, E. Z. A Generative Model for the Joint Registration of Multiple Point Sets. in Computer Vision – ECCV 2014 (eds. Fleet, D., Pajdla, T., Schiele, B. & Tuytelaars, T.) 109–122 (Springer International Publishing, Cham, 2014).

79. Fischler, M. A. & Bolles, R. C. Random sample consensus: a paradigm for model fitting with applications to image analysis and automated cartography. Commun. ACM 24, 381–395 (1981).

80. Bragulat-Teixidor, H., Hossain, M. J. & Otsuka, S. Visualizing Nuclear Pore Complex Assembly In Situ in Human Cells at Nanometer Resolution by Correlating Live Imaging with Electron Microscopy BT - The Nuclear Pore Complex: Methods and Protocols. in (ed. Goldberg, M. W.) 493–512 (Springer US, New York, NY, 2022). doi:10.1007/978-1-0716-2337-4_31.

81. Mastronarde, D. N. Automated electron microscope tomography using robust prediction of specimen movements. J Struct Biol 152, 36–51 (2005).

82. Eastman, P. et al. OpenMM 7: Rapid development of high performance algorithms for molecular dynamics. PLoS Comput Biol 13, e1005659- (2017).

83. Liu, S., Wang, C., Latham, A. P., Ding, X. & Zhang, B. OpenABC enables flexible, simplified, and efficient GPU accelerated simulations of biomolecular condensates. PLoS Comput Biol 19, e1011442- (2023).

84. Michaud-Agrawal, N., Denning, E. J., Woolf, T. B. & Beckstein, O. MDAnalysis: A toolkit for the analysis of molecular dynamics simulations. J Comput Chem 32, 2319–2327 (2011).

85. Boerner, T. J., Deems, S., Furlani, T. R., Knuth, S. L. & Towns, J. ACCESS: Advancing Innovation: NSF’s Advanced Cyberinfrastructure Coordination Ecosystem: Services & Support. PEARC 2023 - Computing for the common good: Practice and Experience in Advanced Research Computing 173–176 (2023) doi:10.1145/3569951.3597559.

86. Fuertes, G. et al. Decoupling of size and shape fluctuations in heteropolymeric sequences reconciles discrepancies in SAXS vs. FRET measurements. Proceedings of the National Academy of Sciences 114, E6342–E6351 (2017).

87. Mirdita, M. et al. ColabFold: making protein folding accessible to all. Nat Methods 19, 679–682 (2022).

88. Jumper, J. et al. Highly accurate protein structure prediction with AlphaFold. Nature 596, 583–589 (2021).

89. Ladd, A. J. C. & Woodcock, L. V. Triple-point coexistence properties of the lennard-jones system. Chem Phys Lett 51, 155–159 (1977).

90. Varadi, M. et al. AlphaFold Protein Structure Database: massively expanding the structural coverage of protein-sequence space with high-accuracy models. Nucleic Acids Res 50, D439–D444 (2022).

91. Evans, R. et al. Protein complex prediction with AlphaFold-Multimer. bioRxiv 2021.10.04.463034 (2021) doi:10.1101/2021.10.04.463034.

92. Latham, A. P. & Zhang, B. On the stability and layered organization of protein-DNA condensates. Biophys J 121, 1727–1737 (2022).

93. Henderson, J. R. & and Lekner, J. Surface oscillations and the surface thickness of classical and quantum droplets. Mol Phys 36, 781–789 (1978).

94. Benayad, Z., von Bülow, S., Stelzl, L. S. & Hummer, G. Simulation of FUS Protein Condensates with an Adapted Coarse-Grained Model. J Chem Theory Comput 17, 525–537 (2021).

